# TUMOR–PRE-ADIPOCYTE CROSSTALK SUSTAINS BREAST CANCER GROWTH VIA RET SIGNALLING

**DOI:** 10.64898/2026.05.05.722993

**Authors:** Sabrina A. Vallone, Angela Lara Montero, Alba C. Arcones, Irene Ruiz-Garrido, Marcos D. Palavecino, Mercedes A. Montani, Juan I. Jiménez-Loygorri, Ivana Nikolic, Martín García Solá, Luis Leiva-Vega, Marta León, Gladys N. Hermida, Elena Rodríguez, Paula Aguirre, Daniela Maltagliatti, Diego Flaks, Noemi Jordanovski, Eliana Querol, Laura Leguina, Silvia Vornetti, Gabriela Acosta, Eva Wertheimer, Omar A. Coso, Carolina Schere-Levy, Juan P. Fededa, Manuel de la Mata, Edith C. Kordon, Guadalupe Sabio, Albana Gattelli

**Affiliations:** CONICET-Universidad de Buenos Aires, Instituto de Fisiología, Biología Molecular y Neurociencias (IFIBYNE), Buenos Aires, Argentina; Universidad de Buenos Aires (UBA), Facultad de Ciencias Exactas y Naturales (FCEN), Pabellón II, Ciudad Universitaria C1428EGA CABA, Buenos Aires, Argentina; Centro Nacional de Investigaciones Oncológicas (CNIO), Melchor F. Almagro 3 28029, Madrid, Spain; Centro Nacional de Investigaciones Cardiovasculares (CNIC), Melchor F. Almagro 3 28029, Madrid, Spain; Departamento de Biodiversidad y Biología Experimental (DBBE), Biología de Anfibios-Histología Animal, FCEN, UBA, Argentina; CONICET-Instituto de Investigaciones Biotecnológicas (IIB), Campus Miguelete IIB - Av. 25 de mayo 1650 San Martín, Buenos Aires, Argentina; Universidad Nacional de San Martín (UNSAM), Argentina; Hospital Oncológico Marie Curie, Buenos Aires, Argentina; CONICET-Centro de estudios farmacológicos y botánicos (CEFYBO), Buenos Aires, Argentina

## Abstract

Adipose tissue is the dominant stromal component of the breast, yet whether breast tumors exploit adipocyte plasticity to support cancer growth remains unclear. Here, we show that breast tumors actively disturb adipocyte differentiation, generating an immature tumor-adjacent adipose niche enriched in pre-adipocytes that directly promotes tumor progression. In human breast cancer samples, adipocytes located near tumors acquire a pre-adipocyte–like state. Functional studies demonstrate that pre-adipocytes enhance tumor cell proliferation both *in vivo* and *in vitro*. Mechanistically, we identify tumor-intrinsic RET signaling as a key regulator of this interaction. The RET receptor is a clinically relevant target expressed in breast cancer. RET drives a PDGF-B–dependent paracrine program that maintains pre-adipocytes in the tumor milieu. In turn, pre-adipocytes provide RET ligands that reinforce oncogenic signaling in tumor cells. Disruption of the RET-PDGF-B axis limits tumor progression. Together, our findings reveal an active tumor-driven mechanism by which breast tumors regulate adipocyte linage states to sustain growth and identify a novel targetable pathway controlling tumor– adipose tissue communication.

## INTRODUCTION

Tumors remodel their microenvironment, yet whether tumoral cells control lineage commitment in neighboring tissues to support cancer progression remains largely unknown ^1,2^. In the breast, adipose tissue (AT) constitutes the predominant stromal component ^3,4^. AT is a highly plastic organ composed not only of mature adipocytes—cells specialized in lipid storage—but also immune and endothelial cells, fibroblasts and progenitors ^5^. Adipocytes represent a major but still poorly characterized component of the breast tumor microenvironment.

In the setting of breast cancer, tumor epithelial cells are in close anatomical proximity with white AT ^4,6^. Adipocytes and their precursors, the pre-adipocytes, comprise a substantial fraction of the mammary AT in both the normal gland and the cancerous breast ^6^. Pre-adipocytes have been described as immature cells of mesenchymal origin expressing distinct markers ^7,8^. Several studies have reported that adipocytes adjacent to tumors display aberrant morphology, altered metabolism, and transcriptional reprogramming ^9–13^. However, whether these alterations reflect a passive loss of adipocyte identity or an active tumor-driven blocked of adipocyte differentiation remains unclear. Here we show that breast tumors alter adipocyte differentiation leading to accumulation of pre-adipocytes that sustain tumor growth, a process controlled by tumor-intrinsic RET signaling.

RET is a receptor tyrosine kinase that has been involved in several cancers ^14^. Upon RET receptor stimulation by Glial cell line-Derived Neurotrophic Factor (GDNF) family ligands and their co-receptors (GFR∝) ^14,15^, activation of downstream oncogenic signaling pathways such as ERK, contribute to cancer malignancy ^16–18^. Although genomic alterations in RET are rare, RET is overexpressed in approximately 60% of breast tumors and promotes carcinogenesis and therapy resistance ^16,17,19–21^. Using multiple RET-dependent breast cancer models, previous reports have shown that genetic or pharmacological inhibition of RET signaling reduces tumor cell proliferation and metastatic potential ^16,20,22–24^. Consistent with these findings, RET inhibitors have been proposed as a therapeutic approach for solid tumors, including breast cancer ^19,25,26^.

In this study, we show that breast tumors actively suppress adipocyte differentiation, generating an immature tumor-adjacent AT enriched in pre-adipocytes that directly support tumor growth. We further demonstrate that tumor-intrinsic RET signaling orchestrates this process through a PDGF-B-dependent paracrine program that maintains adipocytes in their undifferentiated state. In turn, PAs provide RET ligands, establishing a bidirectional feed-forward signaling loop that sustains oncogenic RET activation and tumor growth.

## RESULTS

### Breast tumor cells disrupt adipocyte differentiation, generating an immature pre-adipocyte-enriched adjacent adipose tissue (AT)

To investigate whether breast tumors are associated with alterations in the surrounding AT, we analysed adipocyte morphology in tumor-adjacent regions from a cohort of breast cancer patients (BC patients). Histological examination of hematoxylin and eosin (H&E)–stained sections revealed a marked reduction in adipocyte size in areas proximal to the tumor (Adjacent AT) compared with more distant regions (Distant AT) of the same breast biopsy (**Fig. 1A, B**), a feature consistently observed across samples and independently of breast tumor molecular subtype (**Fig. S1A; Table S1**; **Table S2**), indicating that it is a general feature of the tumor microenvironment.

**Figure 1.**
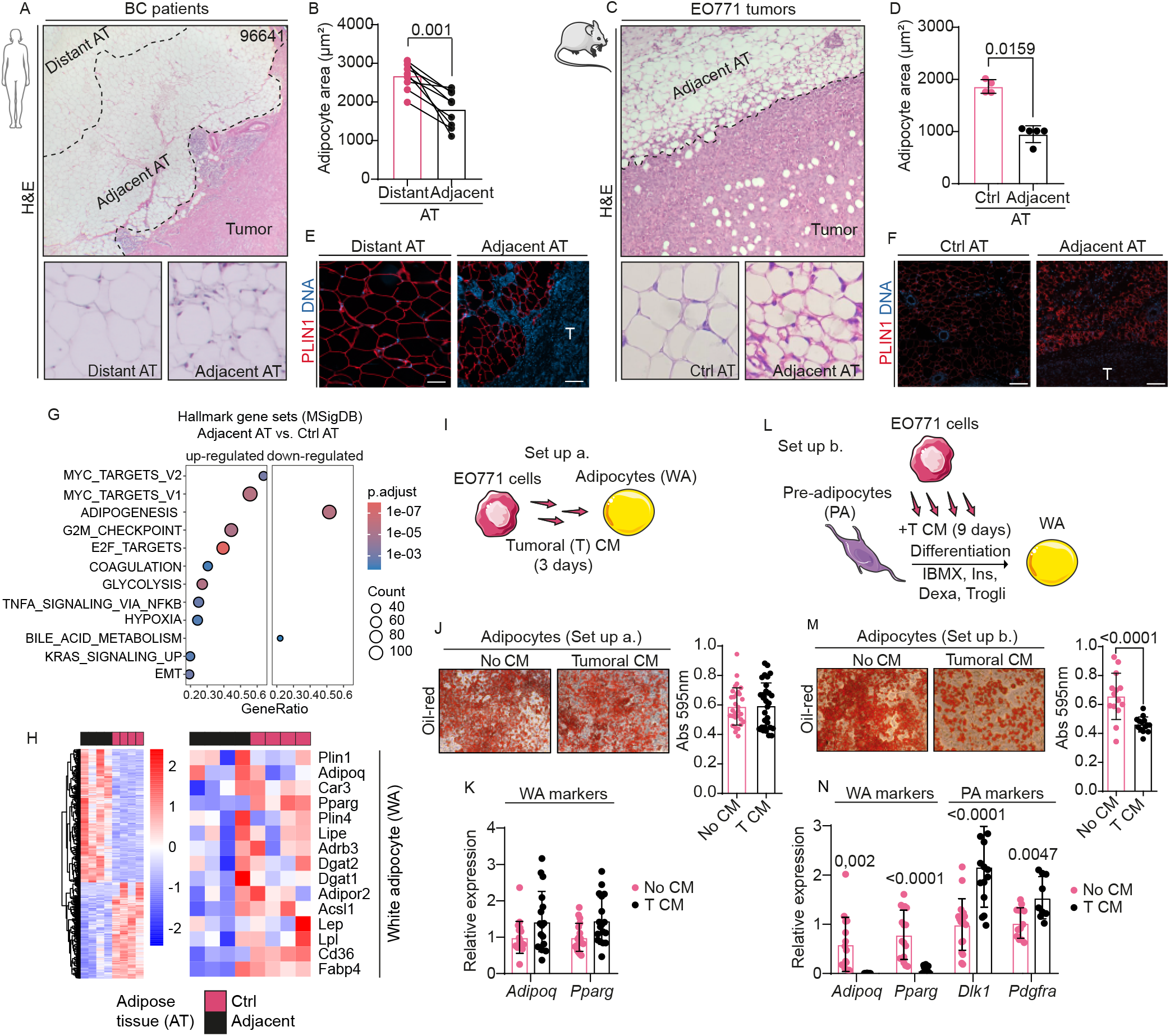
Breast tumor cells are associated with an immature adjacent adipose tissue. (A) H&E-staining shows smaller adipocytes in the proximity of breast tumor in patients. Representative pictures indicating regions of the breast adipose tissue in distant (Distant AT) or adjacent (Adjacent AT) locations to the tumor mass are shown. BC patient ID is indicated on the right corner of the picture. Images were acquired at 10x or 400x magnification. (B) Analysis of average adipocyte area in Distant (>2000 mm) and Adjacent (<2000 mm) regions of the adipose tissue (AT) next to tumor lesion in BC patient biopsies. Each dot represents an individual sample (n=12). Data are presented as means. Unpaired Student’s t test was used; **p=0.001. (C) H&E-staining shows smaller adipocytes in the proximity of mammary tumor in the mouse EO771-tumor model. Representative pictures of either regions of the mammary adjacent adipose tissue (Adjacent AT) or tumor-free adipose tissue (Ctrl AT) are shown. Images were acquired at 10x or 400x magnification. (D) Quantification of adipocyte area in mouse mammary samples from the EO771 model is shown in the graph bars. Each dot represents an individual animal (n=4-5). Data are presented as median ± SD. Mann-Whitney test was used; *p=0.0159. (E-F) Representative images of IF for PLIN1-staining in tumor-adjacent and control areas of adipose tissue in the human breast and mouse model. PLIN1 (red) labels lipid droplet membranes. Nuclei DNA (DNA) were counterstained with DAPI (blue). Images were acquired at 10x magnification. Scale bars=100 μm. (G-H) Hallmark gene set analysis (MSigDB) of RNA-seq data from tumor-adjacent adipose tissue (Adjacent AT) compared with control adipose tissue (Ctrl AT) in the EO771 mouse mammary tumor model from independent animals (n=4). Gene set enrichment significance was assessed using Fisher’s exact test, with p-values corrected for multiple testing using the Benjamini–Hochberg false discovery rate (FDR). Heatmaps show the distribution of gene expression for each experimental group (left), and the expression levels of selected genes representing mature adipocyte markers (right). (I, L) Schematic representation of the experimental set-up of treatments with conditioned medium (CM) after which cultures were subsequently used for analysis. In a Set up a. cultures of terminal differentiated white adipocytes (WA) were treated for 3 days with CM from EO771 cells. In a Set up b. cultures of pre-adipocytes (PAs) under differentiation process were treated for 9 days with CM. (J, M) After treatments lipid content was determined by oil-red staining in the adipocyte cultures and quantified as shown in graphic bars. Representative pictures of oil-red-stained plates are shown for each condition. Images were acquired at 20× magnification. Each dot represents an individual well (Set up a. n=30 from 3 independent experiments, N=3; Set up b. n=14, N=2). Data is presented as mean ± SD. For statistic unpaired Student’s t test was used; corresponding p-values are indicated on bar graphs. (K, N) Either tumoral CM-treated cultures of terminal differentiated WA (Set up a.) or PAs under differentiation process (Set up b.) were used for the analysis of expression of specific adipocyte differentiation markers by RT-qPCR (*Adipoq, Pparg* for WA; *Dlk1, Pdgfra* for PA). Each dot represents an individual well (n=9-18) from 3 independent experiments (N=3). Data is presented as mean ± SD. Unpaired Student’s t or Mann-Whitney tests were used when compared conditions; p-values are indicated on bar graphs. No CM: no CM was added. T CM: Tumoral CM from EO771 cultures. Schemes from experimental procedures along figures were prepared using ServierMedical Art (https://smart.servier.com/), licensed under a creative common Attribution 3.0 unported license.

To assess whether this phenotype is conserved in mice, we analysed AT from female mice bearing orthotopic mammary tumors using the EO771 tumor model ^27^. AT adjacent to tumors (Adjacent AT) displayed reduced adipocyte area compared with AT from tumor-free mammary fat pad (Ctrl AT) (**Fig. 1C, D**). In both human and murine samples, morphological alterations were further confirmed by immunofluorescence staining for Perilipin1 (PLIN1), a membrane marker of mature adipocytes (**Fig. 1E, F**). These changes were spatially restricted to tumor-adjacent regions, suggesting a local effect of the tumor on the surrounding AT.

To determine whether tumor proximity alters AT identity at the molecular level, we performed RNA sequencing (RNA-seq) on anatomically dissected tumor-adjacent AT from the tumor-bearing mice and compared it with AT from the contralateral tumor-free mammary fat pad. Differential expression analysis revealed that peri-tumoral AT (Adjacent AT) presents a completely different transcriptional signature from tumor-free AT (Ctrl AT) (1018 up-regulated genes, 733 down-regulated genes; adjusted p-value <0.05). Tumor-adjacent AT showed decreased activation of adipogenesis pathway and reduced expression of mature adipocyte markers compared to control tissue (**Fig. 1G, H, SI**). These findings extend previous observations describing cancer-associated adipocytes in breast tumors ^28^.

Given that reduced adipocyte size is often associated with impaired adipocyte maturation ^5,10^, we next asked whether tumor cells actively interfere with adipocyte differentiation. To directly test this hypothesis, we assessed the effect of conditioned media derived from tumor cell cultures (CM) on cultured adipocytes ^29^ at different stages of differentiation (**Fig. 1I, L**). Exposure of fully differentiated white adipocytes (WA) to tumor CM did not alter lipid content, as assessed by Oil-Red staining, nor the expression of WA markers, such as *Adipoq* and *Pparg* (**Fig. 1I, J, K**), indicating that tumor-derived signals do not induce dedifferentiation of mature adipocytes under these conditions. In contrast, when tumor CM was added during the differentiation of pre-adipocytes, adipogenic commitment was severely impaired. Cultures exposed to tumor CM showed a marked reduction in lipid accumulation, demonstrating an effective disruption of adipocyte differentiation (**Fig. 1L, M**). This effect was accompanied by reduced expression of key adipogenic regulators and mature adipocytes markers (*Adipoq* and *Pparg*), together with increased expression of markers associated with progenitor states and pre-adipocyte (PA markers), including *Dlk1* and *Pdgfra* (**Fig. 1N**). Consistent with this immature phenotype, DLK1—also known as pre-adipocyte factor 1 (PREF1)— is described as a potent inhibitor of adipogenesis ^7,30^, while PDGFR, the receptor for the Platelet-Derived Growth Factor (PDGF), has been implicated in maintaining pre-adipocytes undifferentiated ^8^.

Together, these findings indicate that tumor-derived signals actively impair adipocyte maturation and promote pre-adipocytes state, providing a mechanistic explanation for the accumulation of pre-adipocyte-like (PA-like) adipose cells observed in tumor-adjacent AT *in vivo*.

### Pre-adipocytes (PA) promote mammary tumor growth and enhance tumor cell proliferation

To evaluate whether pre-adipocytes modulate to tumor *in vivo* behavior, we performed co-injection experiments in which tumor cells were implanted into the mammary fat pad of female mice either alone or together with pre-adipocytes (**Fig. 2A**). Co-injection of tumor cells with pre-adipocytes resulted in increased tumor growth compared with tumors derived from cancer cells alone (**Fig. 2B**). Tumors formed in co-injection with pre-adipocytes displayed a higher proliferative index (**Fig. 2C**), indicating that pre-adipocytes promote tumor expansion *in vivo*.

**Figure 2.**
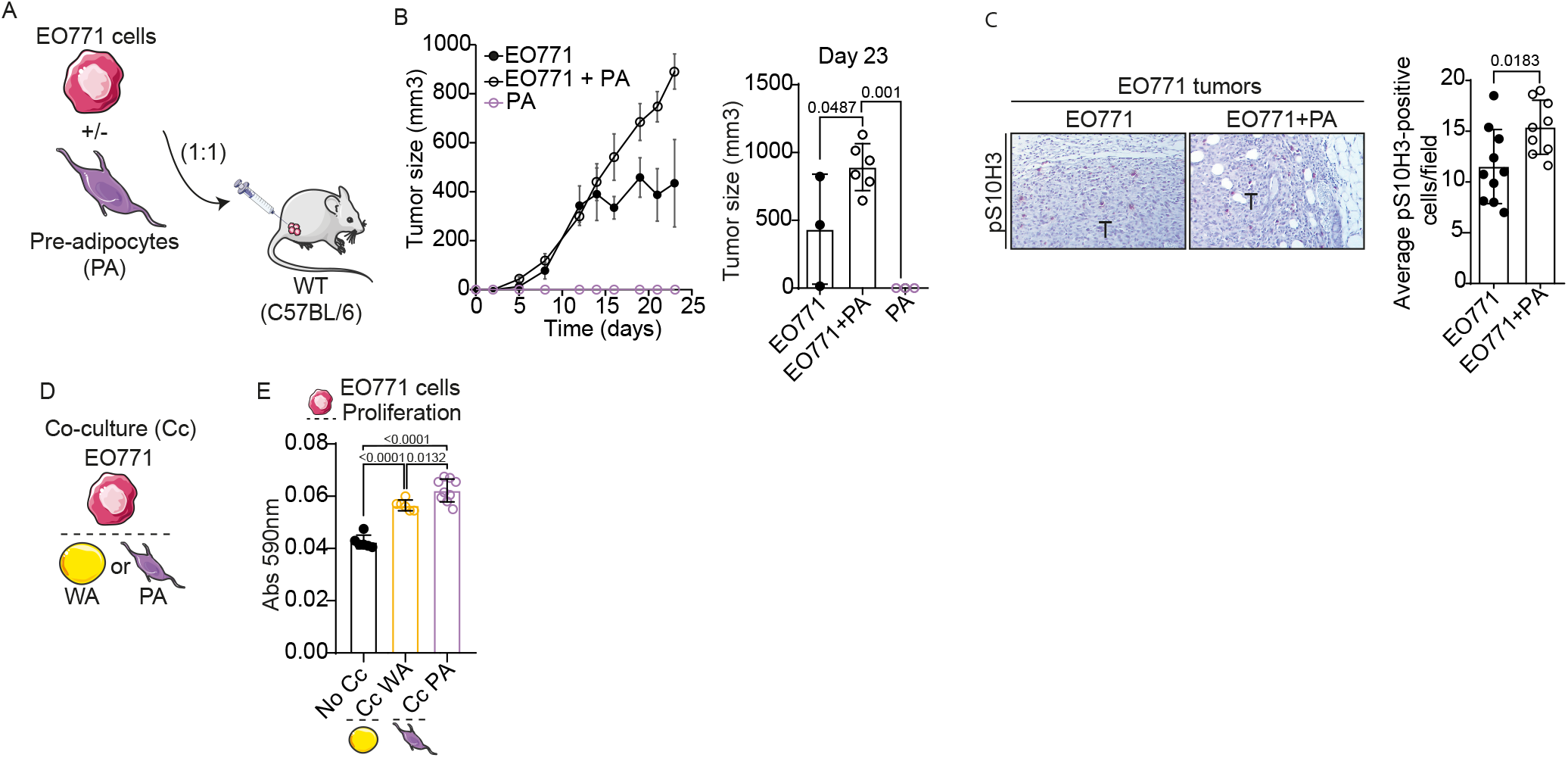
Pre-adipocytes (PAs) increase mammary tumor cell proliferation. (A) Scheme representing the *in vivo* EO771-tumor model where the co-injection of EO771 mammary tumor cells together with PA cells was performed. (B) Representative curves from 2 independent experiments (N=2) showing the tumor growth of the mouse allografts generated by co-injection of both cell types, tumor cell line E0771 and PA cells. Data are presented as mean ± SEM. Each dot in the curve represents the mean (EO771, n=5; EO771+PA, n=6; PA, n=3). Statistics were calculated using One-way ANOVA followed by Tukey’s multiple comparison test at end point (Day 23); p-values are indicated on bar graphics. (C) IHC for a proliferation marker (BrdU) on tumor slides are shown with the corresponding quantification. Data is presented as mean ± SD. Each dot in the curve represents an individual animal (EO771, n=10; EO771+PA, n=9) from 2 independent experiments (N=2). Statistics were calculated using Unpaired Student’s t test; p-values are indicated on bar graphics. Images were acquired at 10× magnification. (D) Schematic representation of the experimental set-up of the transwell co-culture system. Cultures of either differentiated WA or PA cells were co-cultured (Cc) with EO771 tumor cells and subsequently used for the analysis of tumor cell counting. (E) Proliferation was assessed by crystal violet staining followed by absorbance measurement at 590nm after Cc with differentiated WA or PA. Each dot in the plot represents an individual well (n=6-9) from 3 independent experiments (N=3). Statistics were calculated using One-way ANOVA followed by Tukey’s multiple comparison test; p-values are indicated on bar graphics.

We next investigated whether pre-adipocytes directly influence tumor cell proliferation. Using transwell assays, we co-cultured (Cc) tumor cells either with pre-adipocytes (PA) or with differentiated adipocytes (WA) (**Fig. 2D**). These experiments revealed that pre-adipocyte cells enhanced tumor cell proliferation compared with mature adipocytes or control conditions (**Fig. 2E**).

These results indicate that tumor-induced accumulation of pre-adipocytes is not merely a consequence of adipose remodeling but represents a functional stromal mechanism that supports tumor growth.

### Tumor-intrinsic RET signaling regulates mammary AT differentiation

To identify tumor-intrinsic signaling pathways that coordinate tumor progression with AT remodeling, we next investigated candidate oncogenic regulators mediating tumor–adipose crosstalk. Having established that tumor cells actively interfere with adipocyte differentiation and that the resulting accumulation of pre-adipocytes can promote tumor proliferation, we next sought to identify signaling pathways capable of coordinating tumor progression with AT remodeling. In the EO771-tumor mouse model, the transcriptomic analysis of mammary peri-tumoral PA-like AT, identified differentially secreted factors (absolute log2FC >1, *p-value* <0.05; secreted proteins from Human Protein Atlas), among which GDNF, the ligand for the RET oncogene, was prominently induced (**Figure S2A, B, SI**). Receptor tyrosine kinase such a RET and their ligands are key regulators of oncogenic signaling and cellular metabolic adaptation ^31,32^. However, its contribution in mediating mammary tumor-adipose tissue crosstalk remains poorly understood. Notably, analysis of public RNA-seq datasets ^33^ revealed that RET receptor expression is consistently associated with signatures of impaired adipocyte differentiation in the breast tumor-adjacent tissue (**Table S3**). In our patient cohort, the difference in adipocyte area (Δ adipocyte area) positively correlated with higher levels of RET expression in the tumor (**Fig. 3A, B**). These findings prompted us to investigate whether tumor-intrinsic RET signaling activity plays a functional role in tumor–AT crosstalk.

**Figure 3.**
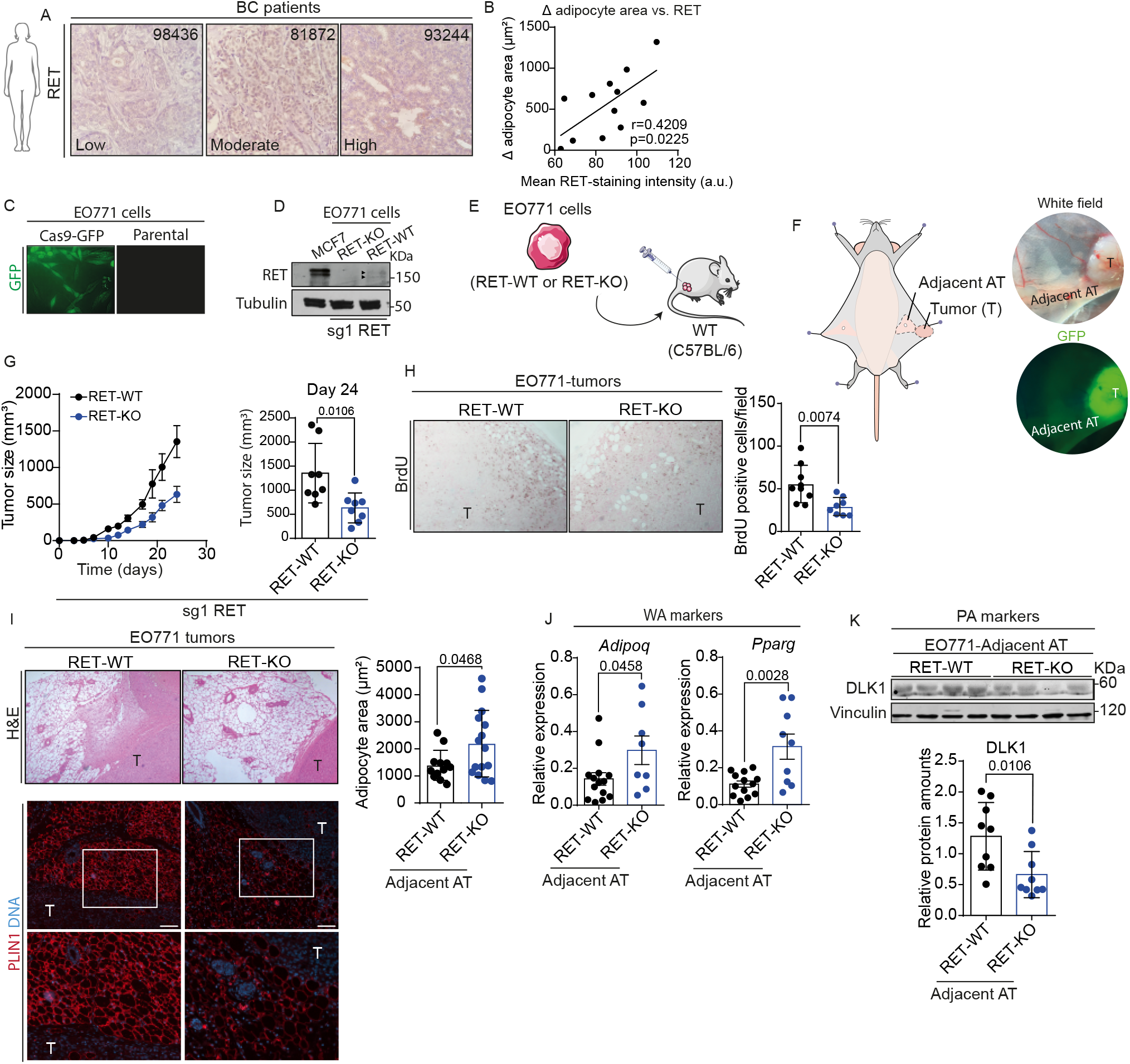
RET-expressing mammary tumors show an immature pre-adipocyte (PA)-enriched adjacent adipose signature. (A) Representative pictures of RET-staining by IHC in breast tumor tissue cores from patients indicating low, moderate and high levels of RET-positive cells. BC patient ID is indicated on the right corner. Images were acquired at 40x magnification. (B) Higher levels of RET expression within tumors are positively correlated with grater difference in the size of adjacent adipocytes in the BC patient cohort. Correlation analysis of the difference in median adipocyte area (D adipocyte area) and tumoral RET expression (by intensity of RET-positive cells in pixels) is shown. Each dot represents an individual sample (n=12). Pearson correlation test was used; *p=0.0225. (C) Representative images showing GFP-driven fluorescence (GFP) in EO771 cultures Cas9-expressing (Cas9-GFP cassette) cells respect to control (Parental) are shown. Images were acquired at 400x magnification. (D) EO771 RET-WT and RET-KO mammary tumor cells (sg1 RET) were analyzed for RET protein expression by WB with specific antibodies, confirming the absence of RET expression in the RET-KO cells. A lysate from MCF7 human breast cancer cells was added as positive control of RET expression. (E) Scheme of RET-KO EO771-tumor allograft model. In this case, RET-WT or RET-KO EO771 tumor cell lines (sg1 RET) were injected into female mice to generate EO771-derived tumors. (F) Scheme showing a dissection and anatomical location of either tumor (T) or tumor adjacent adipose tissue (Adjacent AT) used for posterior analysis. Fluorescence images (GFP) confirmed GFP-positive tumor cells forming the tumor mass. Images were acquired at 1x magnification. (G) *In vivo* tumor growth of RET-WT or RET-KO EO771-tumor allografts (sg1RET). Tumor volumes were measured after mammary fat pad injection of EO771 into female mice (RET-WT, n=8; RET-KO, n=8). Data are presented as mean ± SEM. Each dot in the curve represents the mean of a group of animals from a representative experiment from a total of 8 independent experiments (N=8). In the bar chart, each dot represents an animal. Statistics were calculated using two-tailed unpaired Student’s t test at end point (Day 24); *p=0.0106. (H) BrdU incorporation followed by specific staining were performed on tumor tissue from the EO771-tumor model from 2 independent *in vivo* experiments (N=2). Representative pictures are shown with the indicated quantifications in the graph bars. Each dot represents an individual animal (n=8). Data is presented as mean ± SD. Unpaired Student’s t test was used; **p=0.0074. Images were acquired at 10× magnification. (I) H&E- and PLIN1-staining were performed on tumor and adjacent tissue from the EO771-tumor model. Representative pictures are shown with the indicated quantifications of the adipocyte area in the graph bars. Each dot represents an individual animal (n=16, N=8). Data are presented as median ± SD. Unpaired Student’s t test was used; *p=0.0468. Images were acquired at 10× magnification. Scale bars=100 μm. (J) Analysis of expression of specific adipocyte differentiation markers (*Adipoq, Pparg*, for WA) were performed by RT-qPCR on samples of Adjacent AT from EO771 RET-WT or RET-KO EO771-tumor bearing animals. Each dot represents an individual animal (n=5-21, N=4). Data is presented as mean ± SEM. Unpaired Student’s t or Mann-Whitney tests were used when compared conditions; p-values are indicated on bar graphics. (K) WB analysis showing the expression for DLK1 at protein level on samples of Adjacent AT from EO771 RET-WT or RET-KO EO771-tumor bearing animals. Corresponding quantification is shown. Each dot represents an individual animal (n=9, N=2). Unpaired Student’s t test was used; *p=0.0106.

To address this question, we examined the impact of RET loss on tumor cells and their growth *in vivo* (**Fig. 3C, D, E; Fig. S2C, D, E, F, G**). In this case, for each female mouse, anatomical dissection of both tumor tissue and tumor-adjacent AT was performed (**Fig. 3F**). As expected, orthotopic mammary tumors generated from RET-deficient cells (RET-KO) exhibited reduced growth compared with tumors derived from control cells (RET-WT) (**Fig. 3G, H**), confirming that tumor-intrinsic RET signaling contributes to mammary tumor progression ^16,20^.

Strikingly, reduced growth of RET-deficient tumors was accompanied by marked changes in the AT. Adjacent AT that surround RET-deficient tumors displayed increased adipocyte size compared with RET-WT counterpart (**Fig. 3I**). Furthermore, higher expression of WA adipocyte differentiation markers, including *Adipoq* and *Pparg* were observed in adjacent AT from RET-KO tumors, together with markedly reduced expression of the PA marker DLK1 at the protein level (**Fig. 3J, K, Fig. S3A, B**). Consistent with these results, transcriptomic analysis confirmed that gene programs related to adipogenesis and fatty acid metabolism, which were diminished in AT adjacent to RET-WT tumors, were partially restored in AT adjacent to RET-deficient tumors (**Fig. S3C, D SI**).

These findings indicate that loss of tumor-intrinsic RET signaling is associated with a shift toward a more differentiated adipose tissue phenotype in the tumor microenvironment, thereby identifying RET as a central regulator of adipose tissue differentiation in breast tumors.

### RET signaling drives a PDGF-B–dependent paracrine program conserved across experimental breast cancer models and human breast tumors

To identify paracrine factors induced by RET signaling that could mediate communication with the adipose compartment, we examined transcriptional programs associated with RET activation *in vivo*. For this, we took advantage of previously generated RNA-seq data from in mammary tissue of doxycycline (DOX)-inducible RET/MTB transgenic mouse ^19,20^. In the mammary epithelial hyperplasia driven by constitutively active RET, transcriptomic analysis identified a limited set of RET-responsive secreted factors, among which the PDGF family member *Pdgfa* was prominently induced (**Fig. 4A; Fig. S4A, SI**).

**Figure 4.**
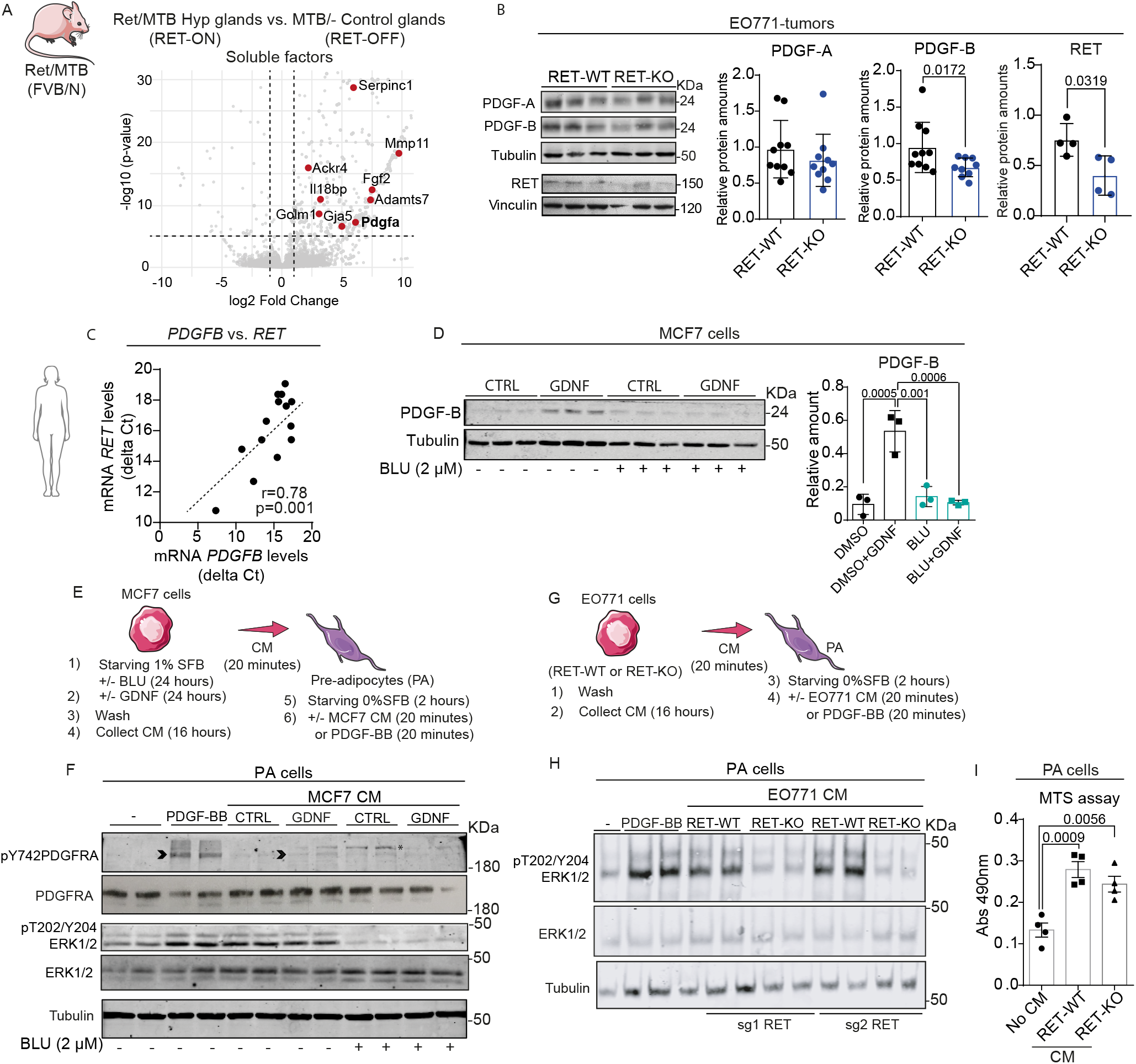
PDGF-B is a RET-driven downstream signaling factor in breast tumors and tumor cells which act on pre-adipocytes (PA) cells. (A) Volcano plot analysis showing the genes encoding soluble factors which are significantly up-regulated in the RET-expressing RET/MTB transgenic glands. p-value <0.01 were considered and calculated by Two-sample t test and log2FC cutoff ± 1. (B) PDGFs levels were confirmed in tumor tissue from the EO771-tumor mouse model (sg1RET and sg2RET) by WB. WB results are representatives of at least three independent animals. Corresponding quantification is shown. Each dot represents an individual animal (n=9-10, 2 independent experiments). Data is presented as mean ± SD. Unpaired Student’s t test was used; *p=0.0172. As control, RET levels in lysates of EO771-tumor tissue (n=4) were analyzed by WB. (C) Correlation analysis of *PDGFB* expression and tumoral *RET* in tumor tissue patient samples from the new cohort. Each dot represents an individual sample (n=14). Pearson correlation test was used; **p=0.001. (D) MCF7 cultures were stimulated with GDNF (25 ng/ml) in absence (DMSO) or presence of RET inhibitor BLU-667 (BLU, 2mM), lysates were obtained and analyzed for PDGF ligands by WB. Results are representatives of at least 3 independent experiments. Corresponding quantification is shown. Each dot represents an individual well (n=3). Data is presented as mean ± SD. One-way ANOVA test was used followed by Tukey’s multiple comparation test p-values are indicated on bar graphics. (E) Scheme of breast cancer cells conditioned medium (CM) treatments on pre-adipocyte (PA). CM from MCF7 (MCF7 CM) previously stimulated with GDNF (25 ng/ml) after treatments with RET inhibitor (BLU, 2mM) was used to treat PA cultures. As control of PDGFR activation, a condition with PDGF-B ligand (PDGF-BB, 0,1 mg/ml) stimulation was performed. (F) Levels of activation of PDGFRA and ERK measured by WB in the PA-treated cultures as indicated. Duplicates are shown in the blots. Arrowhead indicates the correct band. *: unspecific band. (G) A scheme showing experiments where PA cultures were treated with CM from EO771 RET-WT or RET-KO tumor cells (EO771 CM). (H) Levels of activation of ERK were measured (including a time course) by WB in the PA-treated cultures as indicated. The results are shown for two independent EO771 RET-KO cell lines. (I) MTS assays were performed to assess changes in viability in PA-treated cultures with EO771 RET-WT or RET-KO cells CM (48 hours). Each dot represents an individual well (n=4). Data is presented as mean ± SD. One-way ANOVA test was used followed by Tukey’s multiple comparation test; p-values are indicated on bar graphics.

In contrast, PDGF-B, rather than PDGF-A, was consistently associated with RET expression in mammary tumor tissue across *in vivo* mouse models (**Fig. 4B; Fig. S4B**). In human breast tumors, RET expression positively correlated with *PDGFB* levels, with higher *PDGFB* expression in RET-high compared with RET-low tumors (**Fig. 4C; Table S4**). *In vitro*, pharmacological inhibition of RET by BLU-677 (BLU) ^34^ in RET-expressing ^23^ MCF7 and T47D human breast cancer cell lines blocks RET-ligand (GDNF)-induced PDGF-B expression (**Fig. 4D, Fig. S4C**), establishing PDGF-B as a downstream effector of RET signaling in breast tumor cells. Together, these results demonstrate that RET signaling activates a conserved PDGF-B– dependent paracrine program in experimental mouse mammary tumor models and human breast cancer.

To directly test whether tumor-intrinsic RET signaling controls activation of PDGF signaling in the adipose compartment, we exposed PA cultures to conditioned media from RET-proficient or RET-impaired tumor cells. CM from RET-activated tumor cells (MCF7 CM) induced phosphorylation of PDGFRα and downstream activation of ERK signaling in pre-adipocytes (**Fig. 4E, F**), whereas media derived from RET-inhibited or RET-deficient tumor cells (EO771 CM) failed to activate this pathway (**Fig. 4 E, F, G, H**). These results demonstrate that tumor-intrinsic RET signaling is required for the release of soluble factors that activate PDGFRα signaling in pre-adipocytes. Consistently, RET-dependent secreted factors in CM enhanced pre-adipocytes viability (**Fig. 4 I**), an effect reproduced by recombinant PDGF-BB and prevented by pharmacological inhibition of PDGFR signaling (**Fig. S4D, E**). These data indicated that tumor-intrinsic RET signaling promotes a PDGF-B-dependent paracrine mechanism that sustains pre-adipocytes survival.

### Tumor-promoting activity of pre-adipocytes requires RET signaling in cancer cells and provides RET ligands

Having established that pre-adipocytes promote mammary tumor growth and that tumor-intrinsic RET signaling disturbs AT differentiation, we next asked whether the tumor-promoting activity of these pre-adipocytes depends on RET signaling in tumor cells. To address this, we analysed the effect of pre-adipocyte co-injection on tumor growth in the context of RET deficiency (**Fig. 5A**). Crucially, co-injection of pre-adipocytes enhanced the growth of RET-proficient tumors (RET-WT) but failed to increase the growth of RET-deficient tumors (RET-KO), indicating that the pro-tumoral effect of PA requires RET activity in tumor cells (**Fig. 5B**). These results show that pre-adipocytes promote tumor growth in a RET-dependent manner, identifying RET as a key mediator of tumor–PA interactions.

**Figure 5.**
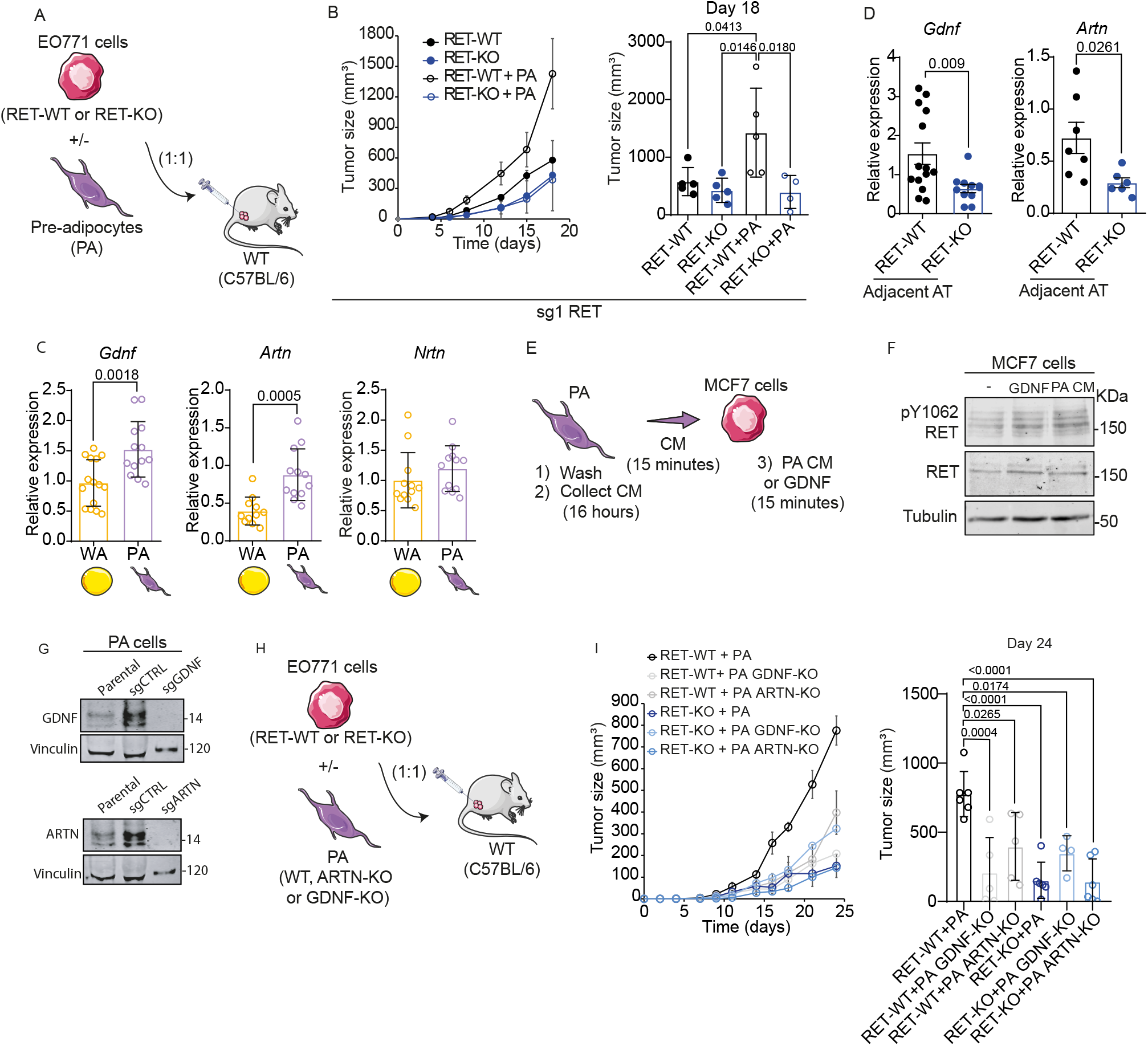
Pre-adipocyte (PA) cells increase tumor growth in a RET-dependent manner through RET ligands production. (A) Scheme representing the *in vivo* experiment of co-injection of EO771 RET-WT vs RET-KO together with PA cells. (B) Representative curves of tumor growth of the mouse allografts generated by co-injection of both cell types, tumor cell lines (sg1RET) and PA cells, are shown (N=2). Data are presented as mean ± SEM. Each dot in the curve represents the mean (RET-WT, n=5; RET-KO, n=5; RET-WT+PA, n=5; RET-KO+PA, n=5; PA, n=3). In the bar chart, each dot represents an animal. Statistics were calculated using One-way ANOVA followed by Tukey’s multiple comparison test at end point (Day 18); p-values are indicated on bar graphics. (C) Expression levels of RET ligands (*Gdnf, Artn* and *Nrtn*) were analyzed in PA cell cultures versus differentiated WA. Each dot represents an individual sample (n=12-14). Data are presented as mean ± SD. Student’s test was used; p=**0.0018, p=***0.0005 (D) Expression levels of RET ligands (*Gdnf* and *Artn*) were analyzed in Adjacent AT from RET-WT or RET-KO EO771-tumors from the allograft model. Each dot represents an individual animal (n=6-14) from 3 independent experiments. Data are presented as mean ± SEM. Unpaired Student’s t or Mann-Whitney tests were used when comparing conditions; p-values are indicated on bar graphics. (E) Scheme of PA cells conditioned medium (PA CM) treatments on MCF7 breast tumor cells. As control of RET activation, GDNF ligand (10 ng/ml) stimulation was performed. (F) Levels of activation of RET were analyzed by WB on MCF7-treated cells as indicated. WB results represent 2 independent experiments (N=2). (G) PA expressing RET ligands (GDNF- or ARTN-WT) and GDNF- or ARTN-KO PA cells were analyzed for ligand expression by WB, confirming the absence of RET ligand expression in the KO cells. (H) Scheme representing the *in vivo* experiment of co-injection of PA cells GDNF/ARTN-WT or GDNF/ARTN-KO with EO771 RET-WT or RET-KO vs together with PA cells. (I) Representative curves of tumor growth of the mouse allografts generated by co-injection of both cell types, tumor cell lines and PA cells, are shown. Data are presented as mean ± SEM. Each dot in the curve represents the mean (RET-WT + PA, n=6; RET-KO + PA, n=4; RET-WT + PA GDNF-KO, n=5; RET-WT + PA ARTN-KO, n=5; RET-KO + PA GDNF-KO, n=4; RET-KO + PA ARTN-KO, n=6). In the bar chart, each dot represents an animal. Statistics were calculated using One-way ANOVA followed by Tukey’s multiple comparation test; p-values are indicated on bar graphics.

We then investigated whether the AT compartment contributes to tumor growth by providing ligands capable of activating RET in cancer cells. RET receptor is more potently stimulated by its ligand GDNF but can be additionally activated by other family members (NRTN, ARTN, PSPN) ^35^. Isolated pre-adipocyte cells (PA) express higher levels of RET-ligands *Gdnf* and *Artn* compared with differentiated adipocytes (WA) (**Fig. 5C**), identifying the immature adipose compartment as a major source of RET ligands. Notably, GDNF expression was not altered by PDGF-B/PDGFR signaling in pre-adipocytes (**Fig. S5A**), suggesting that RET ligand production reflects intrinsic characteristics of the immature adipose state. In addition, analysis of published single-cell RNA-seq datasets from human mammary normal tissue as well as breast tumor-adjacent tissue, shows co-expression of *Gdnf* with the specific pre-adipocyte marker (*Dlk1*) (**Fig. S5B, C**). In the mouse model, RET ligands expression is lower in the AT adjacent to RET-deficient tumors (RET-KO) compared with AT from RET-WT tumors (**Fig. 5D**). Consistently with these findings, conditioned media from pre-adipocytes (PA CM) induced RET phosphorylation in tumor cells comparable to that observed upon stimulation with recombinant GDNF, demonstrating that pre-adipocytes–derived soluble factors are sufficient to activate RET signaling (**Fig. 5E, F**).

To assess the functional relevance of PA-derived RET ligands, we genetically depleted either GDNF or ARTN in PA cells (**Fig. 5G**). Loss of RET ligands in PAs (PA GDNF-KO or PA ARTN-KO) markedly reduced their ability to promote tumor growth *in vivo*, whereas depletion had no effect on RET-deficient tumors (**Fig. 5H, I**).

Together, these findings establish a bidirectional feed-forward loop in which tumor-intrinsic RET signaling maintains an immature adipose niche that, in turn, sustains oncogenic RET activation and tumor growth. Thus, this work positions mammary pre-adipocytes as an active component of the tumor microenvironment that directly influences breast cancer cell proliferation and represents a therapeutic target warranting further exploration.

## DISCUSSION

Adipose stroma consists of several distinct cell types, not just fat-storing adipocytes, also compressing other stromal cells and progenitors such as pre-adipocytes ^5^. Given that AT is a major stromal component of the breast organ, interactions between breast cells and adipose-resident populations are likely to influence breast tumor behavior. In normal mammary AT, pre-adipocytes represent a minor stromal population ^6^ and their contribution to breast cancer progression has remained unclear. In this work, we show that breast tumors actively remodel AT by enforcing a progenitor-like state in tumor-adjacent adipocytes, thereby generating a stromal niche that directly supports tumor growth.

Remodeling of AT is a common feature of tumors and reports have showed that adipocytes adjacent to tumors, commonly referred as cancer-associated adipocytes (CAAs) ^9^, display altered features. In this context, CAAs have been identified at the invasive front of breast tumors and other adipose-rich cancers ^36–39^. Direct contact between tumor cells and adipocytes is reported to generate spindle-like cell states that contribute to the cancer-associated fibroblast (CAF) compartment rather than to differentiated adipocytes ^9,40^, complicating the definition of adipose-derived stromal populations in tumors. In addition, lineage tracing shows that breast adipocytes might de-differentiated into a variety of mesenchymal cell states like pre-adipocytes ^11^ extending the concept of adipose remodeling beyond canonical CAAs.

Here, we characterized the association of breast tumors with an immature adjacent AT: adipocytes close to the tumor are smaller and display altered expression of differentiation markers, while distant AT remains largely unaffected. This phenotype is reproduced in mouse models, indicating that it is conserved among species and imposed locally by the tumor.

Tumor-associated AT immaturity has been linked to both mechanical compression and tumor invasion in breast cancer models ^38,41,42^, consistent with the localized adipose remodeling observed in RET-positive tumors, our results show that tumor-derived signals selectively interfere with adipocyte differentiation at the pre-adipocyte stage explaining the accumulation of PA-like cells in tumor-adjacent AT *in vivo*. We further demonstrate that PAs actively promote tumor cell proliferation and tumor growth, and that supportive effect depend on RET signaling in tumor cells. As with other stromal populations such as fibroblasts (including CAFs) and endothelial cells, PAs express PDGFR ^28,43^, but they can be distinguished by DLK1 expression ^7^, which we find co-localizes with RET ligand expression in tumor-associated AT. Thus, PAs emerge not merely as passive stromal components but as active regulators of tumor progression within the adipose niche.

At the mechanistic level, we reveal PDGF-B as a RET-dependent tumor-derived factor that acts on PDGFR-positive adipose cells, promoting maintenance and survival of the pre-adipocyte population. Among the soluble mediators implicated in tumor-stroma interactions, PDGF-B emerges as a strong candidate linking tumor cells and AT, as its expression positively correlates with RET in both human and mouse tissues. We demonstrate a paracrine signaling axis through which tumor-intrinsic oncogenic signaling directly shapes stromal cell fate. In parallel, we show that PAs are a local source of RET ligands, including GDNF and ARTN, which are enriched in tumor-adjacent AT. While GDNF is the primary RET activator ^35^, ARTN has been implicated in resistance and malignancy of tumors ^44,45^. We tested genetic ablation of these RET ligands in PAs and found that both reduce RET-driven tumor growth, demonstrating a functional role for adipose-derived RET signaling. Together, these findings define a feed-forward signaling loop in which RET-positive tumor cells maintain an immature adipose niche that, in turn, sustains RET activation and tumor growth.

Independently of tumor subtype, RET is overexpressed in breast cancer and despite the low prevalence of activating mutations, clinical trials are testing RET inhibitors ^46^. Our findings indicate that RET oncogenic signaling functions as a mediator of tumor–AT communication. Targeting this tumor–stroma signaling axis may therefore represent a strategy to simultaneously disrupt oncogenic signaling and beneficially remodel the tumor microenvironment.

## Supporting information

Supplementary figures & legends (Fig. S1-5) & Supplementary tables & footnotes (Table S1-4)

## RESOURCE AVAILABILITY

### Lead Contact

Correspondence should be addressed to

### Materials availability

Correspondence and requests for materials should be addressed to

### Data and code availability

The RNA-sequencing data have been submitted to the Gene Expression Omnibus (GEO) and assigned to the identifier GSE324352 and the previously published GSE83897 ^20^.

## EXPERIMENTAL PROCEDURES

### Ethics statement: human cancer tissue source and animal experimentation

This research complies with all relevant ethical regulations. The human sample collection and preparation were reviewed and approved by the Marie Curie Municipal Oncology Hospital Research Ethics Committee (Protocol Reference No: 360; Dated: 12/06/2015). The cohort includes female breast cancer patients who underwent surgery between 2016 and 2020 with all subjects providing written informed consent. Data was collected on demographic information and coexisting medical conditions (e.g., age, gender, histological grade, etc.). This research was conducted according to the principles of the Declaration of Helsinki ^47^.

All animal procedures were conformed to EU Directive 86/609/EEC and Recommendation 2007/526/EC regarding the protection of animals used for experimental and other scientific purposes, enacted under Spanish law Real Decreto 53/2013 and Argentinian guidelines. All experimental protocols (PE) implemented on mice were reviewed and approved by the review board on the use of living animals by the corresponding Institutional Animal Care and Use Committee (IACUC) at Exact and Natural Sciences Faculty (FCEN) from University of Buenos Aires (UBA), Argentina; and at the National Center of Oncology Investigation (CNIO), Spain (IACUC protocols PE-53 and PE-116; PROEX 289.2/23). The maximal tumor burden (10% of the mouse body mass) was permitted by IACUC, and all experiments involving mouse xenografts were terminated before the tumor size reached the maximal tumor volume allowed. FVB/N transgenic (originally from Taconic) and C57BL/6 wild-type (originally from Jackson) mice were purchased from the Animal House at FCEN, UBA (CABA, Argentina) and maintained in a specific pathogen-free facility. In some experiments C57BL/6 mice were purchased from the Animal facility at CNIO (Madrid, Spain).

### Adipocyte area quantification

Adipocyte size and number were quantified using the AdipoSoft plugin implemented in ImageJ/Fiji. Hematoxylin and eosin (H&E)–stained histological sections were imaged using an Olympus CX31 microscope at X10 magnification. Images were analyzed using predefined parameters optimized for adipose tissue detection, including minimum and maximum cell area, circularity constraints, and contrast thresholds. The plugin automatically identified individual adipocytes based on membrane boundaries and segmented them into discrete regions of interest. Detected cells were visually inspected to verify correct segmentation, and misidentified or incomplete cells were excluded when necessary. For each image, AdipoSoft generated quantitative outputs including adipocyte area, diameter, and cell count, which were exported for subsequent statistical analysis. The values were initially obtained in pixels and subsequently converted to micrometers (µm) based on image calibration. For each sample, 3-6 regions were analyzed and mean, and median adipocyte areas were calculated. The Δ adipocyte area was defined as the difference in the median of adipocyte area in the AT between distant (Distant AT) and adjacent (Adjacent AT) locations. For mouse mammary tissues from the EO771-tumor model, samples were processed, and adipocyte area was analyzed and calculated as described before.

### Cell lines and cell culture reagents

Breast cancer RET-expressing breast tumor cell lines, MCF7 (#HTB-22, ATCC) and T47D (obtained from Dr. NE. Hynes laboratory) of human origin as well as EO771 (#C0006048, AddexBio) of murine origin, were used along with this work. MCF7, T47D, and EO771 cells were cultured in high-glucose DMEM (Gibco) supplemented with 10% fetal bovine serum (FBS; Internegocios) and 1% penicillin/streptomycin (Gibco). A pre-adipocyte (PA) cell line, developed in the laboratory of Dr. G. Sabio, was previously generated via immortalization of primary murine mammary pre-adipocyte cultures ^29^. PA cells were maintained in DMEM/F12 containing 8% FBS and 1% penicillin/streptomycin and retain the capacity to differentiate into mature adipocytes, as detailed later. HEK-293T cells (obtained from Dr. de la Mata laboratory) used for lentiviral vector production were cultured in DMEM/F12 with 10% FBS and 1% penicillin/streptomycin (Gibco). Cell lines were used with no more than 20 passages.

Culture media were supplemented with 2 mmol/l L-glutamine (#A2916801, Gibco). Cells were monthly tested for mycoplasma contamination by PCR using specific primers (Myco_Fw - 5’-CGCCTGAGTAGTACGTCGC-3’; Myco_Rv - 5’-CGGTGTGTACAAGACCCGA-3’; U4_Fw - 5’-GCCAATGAGGTTTATCCGAGG-3’; U4_Rv - 5’-TCAAAAATTGCCAATGCCG-3’) and cultured in standard tissue culture-treated plasticware under humidified conditions at 37°C and 5% CO_2_.

GDNF, PDGF-BB (#450-10, #315-18, Peprotech) and GFRα1 (#714-GR, R&D) were stored at -80°C and later used on *in vitro* experiments at 10-25 ng/ml, 0.1 mg/ml and 100 ng/ml, respectively, which is specified in the legends. The selective tyrosine kinase inhibitors for RET, BLU-667 (HY-112301, MCE) named BLU, or for PDGFRA, named Imatinib (SML-1027, Sigma-Aldrich) named as IMA, were prepared in DMSO and stored at -20°C. The inhibitors were used at the following concentrations for cell culture work: BLU (2μM), IMA (1μM).

### Adipocytes differentiation cultures

A total of 20.000 pre-adipocyte (PA) cells were seeded in 12-well plates and cultured in complete medium. As previously described ^29^, to differentiate PA cultures into white adipocytes (WA), culture medium was replaced every two days and sequentially supplemented with adipogenic inducers. For that, at 2 days post-seeding (day 1 of differentiation), the medium was replaced with differentiation-inducing medium containing troglitazone (Trogli; #3114/10, Tocris Bioscience), 3-isobutyl-1-methylxanthine (IBMX; #I5879, Sigma), dexamethasone (Dexa; #D1756, Sigma), and insulin (Ins; #I9278, Sigma). On day 3, the medium was replaced with fresh medium containing troglitazone and insulin. On days 5 and 7, the medium was further replaced with medium supplemented only with insulin. By 9 days, cells exhibited adipogenic differentiation, as evidenced by the presence of intracellular lipid droplets clearly visible under light microscopy.

### Lentivirus production and CRISPR/Cas9 Knockout (KO) cell lines generation

To generate a stable CRISPR/Cas9 expression system, a lentiviral vector containing Cas9 coupled to green fluorescent protein (GFP) based on Ruc-GFP (pRRL-U6-gRNA-SFFV-Cas9-2A-GFP) were obtained by NheI digestion and re-ligation (enzymes by NEBiolabs). Lentiviral vectors expressing sgRNAs with resistance to G418 (Neo^r^ cassette) (LN) were generated from Ruc-Tom (pRRL-U6-gRNA-SFFV-Cas9-2A-TOM). Briefly, the Neo^r^ cassette was excised from the pCI-HA vector with HindIII-BstBI, blunt ended and inserted into AgeI-BsrGI sites within vector Ruc-Tom.

Lentiviral particles were subsequently produced in HEK-293T cells by transfection ^48^ with the packaging vectors pMD2.G (#12259, Addgene) and pCMVR8.74 (#22036, Addgene). Supernatants were collected and low titters (MOI<0.5) were used to transduce EO771 cells. Cas9-expression in cells was corroborated by immunofluorescence (IF) for the fluorescent protein in the cultures and by Western blot (WB) on cell lysates using specific antibodies (anti-GFP #11814460001, Roche Diagnostics, 1:1000; anti- β-actin sc-47778, Santa Cuz, 1:2000, anti-HA #11867423001, Roche Diagnostics, 1:1000). To avoid heterogeneity in Cas9 levels, EO771 Cas9-GFP positive cells were enriched with fluorescence-activated single cell sorting (FACS) using a FACSAria II (Becton Dickinson, San Jose, CA). Cell pools with an average 99% GFP-positive cells were achieved after 3 rounds of sorting (Cas9-GFP pool). Individual clones were further generated by the standard limit dilution method. Briefly, serial dilutions of cells were prepared to obtain single cells in individual wells. Individual clones were finally expanded for further experiments and used to generate the knock-out (KO) cell lines.

For RET depletion, two independent sgRNAs (named Sg1 and Sg2) were design against RET’s first exon-(RET_Sg1Fw - 5’-TCTATGGCGTCTACCGTACA-3’ and RET_Sg1Rv - 5’-TGTACGGTAGACGCCATAGA-3’; RET_Sg2Fw - 5’-CATGCCCTACGGGATGCCCC-3’ and RET_Sg2Rv - 5’-GGGGCATCCCGTAGGGCATG-3’) and cloned into the LN vector. To obtain EO771 control (EO771 RET-WT) or EO771 cells with RET loss (EO771 RET-KO), an empty vector or sgRNA-expressing vector was transduced into Cas9-GFP-expressing cells. The lenti-viral particles were added to the cells with polybrene for 6 hours, and after that G418 (300 ug/mL) was added to select positive cell pools or clones produced by limit dilution as described above. To confirm presence of the RET indel mutations, genomic DNA was extracted by standard methods and the genomic region surrounding the CRISPR/Cas9 target site for RET was amplified by PCR with specific primers (RET_CheckPCR Fw - 5’-AAGCAACTGCCTTTACCCTGT-3’ and RET_CheckPCR Rv - 5’-TCGATGTTTTCCCCATCCCCA-3’). Surveyor assay with T7 endonuclease (#M0302, NEBiolabs) was performed according to the manufacturer’s protocol (95ºC 5 minutes, 95-85°C -2°C/second, 85-25°C - 0.1°C/second) and analyzed on a 2% agarose gel. RET depletion was also confirmed at protein level by WB using a specific antibody (#3223, Cell Signaling, 1:1000).

To generate the PA RET-ligands knock out cell lines (PA GDNF-KO and PA ARTN-KO), transient calcium phosphate co-transfection of HEK-293T cells was carried out with LV-Cas9-scramble (pLV/[CRISPR])-hCas9:T2A:Puro-U6-Scramble[gRNA#1]), LV-Cas9-mARTN (pLV[CRISPR]-hCas9:T2A:Neo-U6>mArtn[gRNA#694]) and LV-Cas9-mGDNF (pLV[CRISPR]-hCas9:T2A:Neo-U6>mGdnf[gRNA#1097]) from Vector Builder, together with psPAX2 (Addgene #12260) and pMD2.G (Addgene #12259) packaging plasmids. Supernatants containing the lentiviral particles were collected 48h and 72h after removal of the calcium phosphate precipitate, centrifuged at 700×g at 4°C for 10min. Viruses were resuspended in cold sterile PBS and titrated by RT-qPCR. In this case, WB was used directly to confirm protein depletion (anti-GDNF #A14639, ABclonal, 1:1000; anti-ARTN #A7949, ABclonal,1:1000).

### Conditioned medium (CM) and treatments

To obtain conditioned media (CM), EO771 tumor cells, either RET-WT or RET-KO, or PA cells were seeded (3 × 10^6^) in 150 mm culture dishes to ensure 80% confluence. After 24 hours, monolayers were washed with PBS 1x, and the culture medium was replaced with starvation serum-free medium (DMEM/F12 0% FBS supplemented with 1% penicillin/streptomycin). Cells were incubated for 16 hours (overnight, ON) under standard culture conditions. The CM was collected and filtered through 0.2 µm pore filters to ensure complete removal of cells.

With respect to tumoral CM, we first assayed the effect of CM derived from EO771 tumor cells on both cultures of mature adipocytes (Set up a.) or cultures under the process of differentiation to white adipocytes (Set up b.). For experiments involving mature adipocytes, fully differentiated cells were treated with CM, which was renewed every 24 hours for 3 days. Control adipocytes were kept in maintenance adipocyte culture medium (No CM) which was also renewed every 24 hours. All treatments lasted 72 hours, after which cells were harvested for further analysis. For the differentiation experiments, CM from tumor cells was added on different days of the differentiation protocol (Day 1, Day 3, Day 5, and Day 7) and compared to control medium. To ensure consistency, tumor-CM was used fresh—either collected immediately prior to treatment or harvested in previous days and stored at −20⍰°C until use. From these experiments, RNA was extracted for quantitative reverse transcription PCR (RT-qPCR), and cells were fixed for Oil-Red staining to assess lipid accumulation.

Next, CM derived from EO771, MCF7 or PA cultures, which were used to stimulate signaling pathways, were concentrated using Vivaspin 2 centrifugal concentrators (5 kDa molecular weight cut-off, MWCO) according to the manufacturer’s instructions (GE Healthcare). The concentrated CM was used directly to stimulate recipient cells and subsequently protein lysates were analyzed for signaling phospho- and - total proteins by WB. In the case of the analysis of tumoral CM-treated PAs we also performed experiments using CM from MCF7 cells. For that, 5 × 10^5^ MCF7 cells were seeded in 60mm culture dishes; after 24 hours, monolayers were washed with PBS 1x, and the culture medium was replaced with DMEM supplemented with 1% FBS and 1% penicillin/streptomycin containing BLU (2 μM) or not (DMSO). After 24 hours, GDNF (25ng/ml) was added for an additional 24 hours. Cells were washed with PBS 1x and incubated for 16 hours (ON) under standard culture conditions. The CM was collected, filtered, concentrated, and used as described before to treated (20 minutes) PA cultures. In this case, PA cultures were starved for 2 hours in DMEM 0% FBS supplemented with 1% penicillin/streptomycin containing IMA (1 μM) or not (DMSO), subsequently protein signaling was analyzed by WB.

In addition, we also conversely used CM harvested from PA cultures to treat breast tumor cell cultures. For that, PA CM was obtained as described to treat for 15 minutes MCF7 cells starved (DMEM/F12 0% FBS 1% penicillin/streptomycin) cultures.

When necessary, stimulation by specific RTK ligands, GDNF or PDGF-BB (10-25ng/ml or 0,1 mg/ml, respectively) was carried out on the assayed cultures of tumor cells (e.i. EO771 or MCF7) or PAs.

### Lipids staining by Oil-Red

Adipocytes cultured in 6-well plates were first washed with PBS 1x and subsequently fixed with 4% paraformaldehyde (PFA) for 10 minutes at room temperature. Following fixation, cells were washed once with PBS and twice with distilled water, then incubated with 500⍰μl of propylene glycol for 5 minutes under agitation. A 1⍰ml solution of 5% Oil-Red in PBS (#O0625, Sigma) was added to each well, and the plates were incubated for 2 hours on a shaker. After staining, the Oil-Red solution was removed; cells were washed three times with distilled water and allowed to air dry for optical microscopy imaging. After imaging, the dye retained in the cells was extracted for quantification by absorbance. To this end, 1⍰ml of isopropanol was added to each well, and the plates were agitated until complete dye solubilization. The resulting solutions were transferred in triplicate to a 96-well plate, and absorbance was measured at 520⍰nm using a CLARIOstar Plus microplate reader (BMG Labtech).

### Co-culture (Cc) experiments in transwells and cell proliferation

For transwell-based *in vitro* co-culture (Cc) experiments PA cells were seeded in the lower compartment of 12-well plates (20.000 cells/well) and subjected to WA differentiation protocol as previously described; additionally, two days before completion of differentiation (Day 7): for the other condition, PA cells were directly seeded at a higher density in 12-well plates (40.000 cells/well). In parallel, EO771 tumor cells were seeded (10.000 cells/well) onto 0.4 μm pore-size transwell inserts (#3460, Corning), which prevent direct cell–cell contact while allowing the exchange of soluble factors. On Day 9, inserts were placed into the upper compartment to initiate Cc and were maintained for 48 hours without medium replacement. At the end of the co-culture period, tumor cells were fixed and stained with crystal violet to assess proliferation.

To determine proliferation, tumor cells were washed with PBS and fixed with 4% paraformaldehyde for 10–15⍰min at room temperature. After fixation, inserts were washed twice with PBS and stained with 0.1% (w/v) crystal violet solution prepared in 20% methanol for 20–30⍰min at room temperature. Excess stains were removed by extensive washing with distilled water until background staining was eliminated, and inserts were air-dried. The dye was then solubilized using 10% acetic acid, transferred in triplicate to a 96-well plate, and quantified by measuring absorbance at 570–590⍰nm using a CLARIOstar Plus microplate reader (BMG Labtech).

### MTS assays

MTS tetrazolium salt colorimetric assay (#G3582, Promega) was used to assess cell viability of treated-PA cells. Optimizations were performed to generate calibration curves to determine the optimal seeding density. Cells were seeded into 96-well plates in complete medium and after 24 hours, cells were serum-starved (1% FBS) for up to 72 hours, when 20 μl of MTS reagent was added to each well under low-light conditions. Control wells contained complete medium, and wells without cells served as blanks. Plates were incubated at 37°C, and absorbance at 490 nm was measured (formazan product) at 20, 40, 60, and 80 minutes using a Varioskan LUX microplate reader (Thermo Fisher). Absorbance values were plotted against cell number to generate a linear calibration curve, which was subsequently used to estimate viable cell numbers in experimental conditions. Based on optimization, PA cells were seeded at densities of 2000 cells per well, and the next day, either treated with CM derived from EO771 RET-KO or RET-WT cells for 48 hours or, pre-treated with IMA (1μM) for 2 hours and subsequently stimulated with PDGF-BB (0,1mg/ml) for 48 hours. After this incubation period, cell viability was assessed using the MTS assay procedure.

MTS assay was also used to determine the half-maximal inhibitory concentration (IC_50_) of BLU-667 (BLU) in MCF7 and T47D cells. For this purpose, MTS assays were performed using concentration curves ranging from 10 to 3000⍰nM of the inhibitor, accompanied by corresponding DMSO concentration curves as controls. An initial seeding density of 10,000–15,000 MCF7 or T47D cells per well was selected for assays performed in 96-well plates. After 24 hours of cell seeding, cells were serum-starved and treated with the indicated concentrations of BLU-667. After an additional 24⍰hours, cells were stimulated with the RET ligand GDNF (25⍰ng/ml) to evaluate cell viability in response to the inhibitor in combination with ligand stimulation. IC_50_ values for each cell line were calculated based on MTS absorbance measurements and microscopic observations, defining IC_50_ as the inhibitor concentration at which cell viability was reduced by 50%. The IC_50_ of BLU-667 was determined to be 2000⍰nM for both cell lines.

### Real-time quantitative PCR (RT-qPCR)

Fragments of tumors or tumor-adjacent AT (Adjacent AT) obtained from anatomic dissection of the left inguinal fourth mammary gland were homogenized for 15 seconds using Ultraturrax T25 (Ika Labortechnik Staufen) in Trizol (Invitrogen), according to the instructions. Pieces from intact inguinal #4 tumor-free mammary gland (CTRL AT) were also processed. Then, the total RNA was isolated by using the NucleoSpin RNA kit. For cell cultures, total RNA was extracted directly with the NucleoSpin RNA kit. One microgram of total RNA was used for reverse transcription according to the MultiScribe Reverse Transcriptase protocol (Thermo Fisher). RT-PCR was performed in a Bio-Rad instrument using a standard reaction mix containing DNA polymerase (K1001 Inbio Highway) up to 40 cycles of amplification.

The program was as follows: 94°C for 2 minutes (min), 40 cycles of 94°C for 15 seconds (s), 60°C for 20 s, 72°C for 1 min/Kb, plus a final extension step of 72°C for 4 min. RT-qPCR was performed on the StepOne Plus instrument (Applied Biosystems) using NZYSpeedy qPCR Green Master Mix. For RT-qPCR, calibration curves were performed for each specific primer pair and used in the calculation for quantification. The program was as follows: 95°C for 5 min, 40 cycles of 95°C for 15 s, 60°C for 20 s, 72°C for 20 s, plus standard program for melting curves.

The sequence of primers (Sigma or Macrogen) utilized for mouse (m) or human (h) are listed below:

m_Gdnf Fw 5’- AGGAGGAACTGATCTTTCGATAT -3’

m_Gdnf Rv 5’- TGGCCTACTTTGTCACTTGTT -3’

m_Artn Fw 5’- CCCTAGCTGTTCTAGCCCTG -3’

m_Artn Rv 5’- AGGGTTCTTTCGCTGCACAA -3’

m_Nrtn Fw 5’- GGGCTACACGTCGGATGAG -3’

m_Nrtn Rv 5’- CCAGGTCGTAGATGCGGATG -3’

m_Prsn Fw 5’- GGCAGATAAGCTCTCATTTGGG -3’

m_Prsn Rv 5’- CACAGTCGGCATGAACCAG -3’

m_Adipoq Fw 5’- TGTTCCTCTTATCCTGCCCA -3’

m_Adipoq Rv 5’- CCAACCTGCACAAGTTCCCTT -3’

m_Pparg Fw 5’- TCGCTGATGCACTGCCTATG -3’

m_Pparg Rv 5’- GAGAGGTCCACAGAGCTGATT -3’

m_Lep Fw 5’- TGTTCCTCTTATCCTGCCCA -3’

m_Lep Rv 5’- CCAACCTGCACAAGTTCCCTT -3’

m_Plin1 Fw 5’- ACAGCAGAATATGCCGCCAA -3’

m_Plin1 Rv 5’- GGCTGACTCCTTGTCTGGTG -3’

m_Wisp1 Fw 5’- CAGCACCACTAGAGGAAACGA -3’

m_Wisp1 Rv 5’- CTGGGCACATATCTTACAGCATT -3’

m_Cebpa Fw 5’- CTGCTCTCCTTTCTCAGGGGT -3’

m_Cebpa Rv 5’- GTGTGCAGTGCTATCATCAA -3’

m_Dlk1 Fw 5’- GCGTGATCAATGGTTCTCCCT -3’

m_Dlk1 Rv 5’- ACTGGCGCAGTTGCTCAC -3’

m_Pdgfra Fw 5’- CTGTTGGAGCTTGAGGGAGAG -3’

m_Pdgfra Rv 5’- CCCATAGCTCCTGAGACCTTCT -3’

m_Icam1 Fw 5’- GTGATGCTCAGGTATCCATCCA -3’

m_Icam1 Rv 5’- CACAGTTCTCAAAGCACAGCG -3’

m_Dpp4 Fw 5’- ACCTTGACCATTACAGGAACTCA -3’

m_Dpp4 Rv 5’- CTAGCGATCCCGTGGTCTTC -3’

m_Cd24 Fw 5’- ACTCAGGCCAGGAAACGTCTCT -3’

m_Cd24 Rv 5’- AACAGCCAATTCGAGGTGGAC -3’

m_18S Fw 5’- TGTTCCTCTTATCCTGCCCA-3’

m_18S Rv 5’- CCAACCTGCACAAGTTCCCTT-3’

m_Gapdh Fw 5’- TGAAGCAGGCATCTGAGGG -3’

m_Gapdh Rv 5’- CGAAGGTGGAAGAGTGGGA -3’

h_PDGFB Fw 5’- CCCGGAGTCGGCATGAAT -3’

h_PDGFB Rv 5’- GGCCCCATCTTCCTCTCCG -3’

h_RET Fw 5’- TCATATGTGGCCGAGGAGGCG -3’

h_RET Rv 5’- CAGTCCTGAGGGCAAATGTTGATG -3’

h_18S Fw 5’- GTAACCCGTTGAACCCCATT -3’

h_18S Rv 5’- CCATCCAATCGGTAGTAGCG -3’

### Western blot (WB)

Cells from the monolayer were washed (PBS 1x) and directly lysed by adding cracking buffer (Tris-HCl 50mM, pH 6.8; SDS 2%; Glycerol 10%; Bromophenol blue 0,006%; and β-mercaptoethanol 1%). For mammary tissue, pieces of tumor-adjacent AT (Adjacent AT) or tumors previously store at -80°C, were homogenized for 15 seconds in RIPA buffer (Tris-HCl20 mM, pH=7.4; EDTA 2mM; NaCl 137 mM; Glycerol 10%; SDS 0,1%; Sodium Deoxycholate 0,5%; Triton X100 1%) containing protease inhibitor (Cocktail set I, Calbiochem, San Diego, CA) and phosphatase inhibitor cocktail (NaF 1mM, sodium glycerolphosfate 40 mM y Na3VO4 1 mM), using Ultraturrax T25 (Ika Labortechnik Staufen, Germany). Lysates were cleared by centrifugation at 12,000 g, and the supernatants were collected and stored at −80°C. Before freezing, an aliquot of lysate was diluted at 1:20 in PBS 1× for determination of the protein concentration with a Bradford assay (Coomasie Blue G250 4%; Ethanol 5%; phosphoric acid 8,5%) using BSA as a standard (Sigma). Extracts of total protein (40-60 μg) prepared in cracking buffer, were incubated for 10 minutes at 95ºC to complete process of denaturalization, separated using sodium dodecyl sulfate polyacrylamide gel electrophoresis (SDS-PAGE) and subsequently transferred onto nitrocellulose (GE HealthCare) blotting membrane. A molecular weight marker was added (Page Ruler Plus prestained marker, Pierce; BlueStar Prestained Protein Marker, Nippon Genetics). The membranes were blocked with 5% casein (Sevelty, Nestle) in TBS-T (Tween 0,1% in buffer TBS 1X) for 1 hour at room temperature. Subsequently, the membranes were incubated overnight at 4°C with the primary antibody diluted in 5% casein TBS-T. The following primary antibodies were used for overnight incubations: anti-RET (#3223, Cell Signaling Technology, 1:1000), anti-p-Y1062RET (sc-2052, Santa Cruz Biotechnology, 1:1000), anti-DLK1 (#10636-1-AP, Proteintech, 1:5000), anti-PDGF-A (sc-9974, Santa Cruz, 1:1000), anti-PDGF-B (sc-365805, Santa Cruz, 1:1000), PDGFRA (AF1062, R&D Systems, 1:1000), p-Y742PDGFRA (AF2114, R&D Systems, 1:1000), anti-ERK (#9102, Cell Signaling Technology, 1:5000), anti-p-T202/Y204ERK (#9101, Cell Signaling Technology, 1:5000), anti-HA (#11867423001, Roche Diagnostics, 1:5000), anti-GFP (#11814460001, Roche Diagnostics, 1:5000) and anti-α-tubulin (MS-581-P1, Neomarkers, 1:2000), anti-β-actin (sc-47778, Santa Cruz, 1:2000) or anti-vinculin (sc-73614, Santa Cruz, 1:2000) as housekeeping loading control proteins. The following day, the primary antibodies were removed, and the membranes were washed in PBS-T.

Subsequently, either the secondary horseradish peroxidase (HRP)-conjugated antibody (Amersham, GE Healthcare) anti-rabbit (NA934, 1:10000), anti-mouse (NA931, 1:10000), anti-goat (sc-2020, Santa Cruz, 1:10000) or fluorophore-conjugated antibody (IRDye, LI-COR) anti-rabbit 800 CW (926-32213, 1:10000), anti-mouse 680 RD (926-68072, 1:10000) or anti-goat 800 RD (926-32214, 1:10000) was added and incubated for 1 hour at room temperature. Finally, the membranes were visualized either using the ECL Western Blotting System (Amersham, GE Healthcare) and CL-XPosure films (Thermo Fisher Scientific) or by Odyssey equipment (LI-COR). For protein level quantifications, target protein as well as housekeeping bands on WBs were quantified using ImageJ. Lysates corresponding to samples from independent animals at each condition were used. Results were expressed as ratios, protein relative to housekeeping protein.

#### Allografts models

EO771 parental cells, RET-KO cell lines and control EO771 cells (RET-WT) were tested in female C57BL/6 wild-type (WT) mice, with 5-8 mice per group. To obtain EO771-tumors, 1 × 10^6^ cells in 100 μl of PBS 1X were injected subcutaneously into the left fourth (#4) mammary fad pad of 5–8-week-old female mice. Tumor growth was monitored every 2 days. Tumor volumes were determined according to the formula: length x diameter 2 × π/6. When EO771-tumors reached ∼ 1200mm^3^ (18-24 days after injection) mice were euthanized and autopsies were performed.

For specific experiments of co-injection, EO771 cells (EO771 RET-KO or EO771 RET-WT) were co-injected with PA cells; parental PA or PA RET-ligand KO cell lines or controls were used. Both cell types were cultured independently, and, on the day of injection, single-cell suspensions were prepared. Each mouse received 1 × 10^6^ cells from a previously mixed composition of tumor cells with PA cells (1:1). As a control, suspensions containing only 1 × 10^6^ of PA cells were assayed.

Tumors as well as the anatomically dissected mammary AT adjacent to the tumor (Adjacent AT) were collected at the end of the experiment and samples were stored at -80ºC or 70% EtOH (previously fixed in 4% PFA) for posterior analysis. Mammary AT from tumor-free gland (#4 right) was stored as control tissue. Dissections were performed under a ZEISS SteREO Discovery.V8 stereomicroscope. Bright-field and GFP fluorescence images were acquired at X1 magnification.

#### Histology, Immunohistochemistry (IHC) and Immunofluorescence (IF)

Mouse tissues were dissected, fixed overnight in 4% paraformaldehyde (PFA), and subsequently stored in 70% ethanol until processing. Paraffin processing, embedding, sectioning, and hematoxylin and eosin (H&E)-staining were performed by the histopathological facility at Maimonides University (Argentina). Images were acquired with PrimoStar3 microscopy (Zeiss) and analyzed.

Immunohistochemistry (IHC) were performed by standard protocols, with the following antibodies: anti-BrdU (#AB6326, Abcam, 1:100), anti-pS10H3 (#9701, Cell Signaling, 1:200) and anti-RET (#sc-167, Santa Cruz, 1:50). Heat-induced epitope retrieval was used with 10 mM sodium citrate pH 6.0 for 10 minutes at sub-boiling temperature, blocking was performed for 1 hour in 2.5% BSA (Sigma) and results were revealed by DAB (Vector Dickinson) using Vectastain ABC kit reagent (PK-6101, Vector Laboratories); counterstaining was performed by hematoxylin (Biopack). Images were acquired with a CX31 microscope (Olympus) or PrimoStar3 microscopy (Zeiss) and analyzed. For the quantification of nuclear markers such as BrdU and pS10H3, positive epithelial cell nuclei were counted in 6-8 randomly selected fields at 400X magnification in mammary tumoral tissue sections. Results were expressed as the average number of positive cells per field. For RET-staining on human tissue, quantification was performed by measuring the stained area. For each sample, three representative tumor areas were selected, and the intensity of DAB staining was quantified using ImageJ software. The mean staining intensity was calculated and expressed in arbitrary units (a.u.) for each sample. Representative images were acquired with PrimoStar3 microscopy (Zeiss) using the ZEISS ZEN 3.12 Blue software. Brightness and contrast were adjusted uniformly across the entire image using Adobe Photoshop.

For IF, paraffin sections were fully deparaffinized by sequential incubations in xylene. Sections were then blocked and incubated with a primary anti-Perilipin1 antibody (#3470, Cell Signaling, 1:1000). After washing, sections were incubated with an Alexa Fluor 647–conjugated anti-rabbit secondary antibody (#A-31573, Thermo Fisher Scientific; 1:1000). Nuclear staining was performed using VECTASHIELD mounting medium containing DAPI (#H-1200-10, Vector Laboratories). Fluorescent images were acquired using an LSM 510 confocal microscope (Zeiss). Individual channels were merged, and images were prepared for figure layout using Inkscape.

In all cases, regions of necrosis, folded or damaged tissue, and specimen edges were excluded from analysis, as these areas can occasionally produce artifactual antibody accumulation and false-positive staining.

#### RNA-sequencing data (RNA-seq) and bioinformatic analysis

Mammary gland tissues were collected from DOX-induced (2 months) Ret/MTB and control MTB/− female mice displaying hyperplastic or normal phenotypes, respectively. Total RNA was extracted using the RNeasy Mini Kit (Qiagen) according to the manufacturer’s instructions. RNA quality was assessed prior to library preparation, and sequencing libraries were generated from 200 ng of total RNA using the TruSeq Stranded mRNA Library Preparation Kit (Illumina). Libraries were pooled and sequenced on an Illumina HiSeq 2500 platform. Base calling was performed using Illumina HCS version 2.2.58 with default parameters, and demultiplexing and FASTQ file generation were carried out using bcl2fastq version 1.8.4 (Illumina). Paired-end reads were aligned to the mouse reference genome (mm10) using STAR (v2.7.11b). Gene models were obtained from the TxDb.Mmusculus.UCSC.mm10.known gene annotation, and gene-level read counts were computed in R using the GenomicAlignments framework by quantifying overlaps between aligned fragments and exonic regions grouped by gene in a union based, strand-specific manner. Gene identifiers (Entrez IDs) were mapped to gene symbols and names using org.Mm.eg.db. Differential expression analysis was performed on gene-level count data, and results were visualized using volcano plots constructed in R. Genes were considered differentially expressed based on thresholds of |log2 fold change| > 1 and −log10(p-value) > 5. Poorly annotated genes (e.g. Rik family genes) were excluded from downstream visualization. Volcano plots were annotated to highlight selected genes encoding predicted soluble factors, based on curated gene lists derived from published literature. Gene Ontology (GO) enrichment analysis was performed in R using the clusterProfiler package on differentially expressed genes to identify overrepresented biological processes, molecular functions, and cellular components. GO annotations were obtained from NCBI, UniProt, and the Gene Ontology Consortium. Enrichment significance was assessed using Fisher’s exact test, and p-values were corrected for multiple testing using the Benjamini–Hochberg false discovery rate (FDR). Pathway enrichment analysis was additionally performed using KEGG annotations.

For distant or tumor-adjacent AT sequencing, total RNA samples (300ng) were converted into sequencing libraries using “QuantSeq 3’mRNA-Seq V2 Library Prep Kit (FWD) for Illumina” (Lexogen, Cat.No. 191). Briefly, library generation is initiated by reverse transcription with oligodT priming, followed by a random-primed second strand synthesis. A UMI Second Strand Synthesis module was used, in which random primers featuring a 6 nt long Unique Molecular Identifier tag is used to address bias correction during downstream data analysis by facilitating the removal of PCR duplicates. Directional cDNA libraries, stranded in the sense of orientation, were completed by PCR with Unique Dual Index primers and sequenced in a single read format on an Illumina instrument (Illumina NovaSeq X) with NovaSeq X Series Reagent Kits. Approximately 8-12 million reads were obtained per sample, and FASTQ file quality control was performed using Fastq-screen (v0.15.2) and MultiQC (v1.18). Reads were trimmed with cutadapt using known, common UMIs and adapters, and reads were aligned and quantified using START (v2.7.11) to the mouse reference genome (GRCm39 mm10). Differential expression analysis was run using DESeq2 (v1.46.0), normalized counts; shrunken log2-FC and adjusted p-values were used for downstream analysis and visualization. Principal components analysis was performed using factoextra (v1.0.7) and FactoMineR (v2.13). Gene-set enrichment was performed using ClusterProfiler (v4.18.4) against HALLMARK and GO databases (MSigDB). Plots were generated using ggplot2 (v4.0.2), ggthemes (v5.2.0), enrichplot (v1.30.4) and pheatmap (v1.0.13). Predicted secreted proteins were obtained from the Human Protein Atlas, then filtered in the dataset based on fold-change (log2FC >1; log2FC <-1) and significance (pvalue <0.05).

Single cell RNA-seq data of published reports from human samples of normal breast or breast tumors ^49,50^ were explored and visualized using the Broad Institute Single Cell Portal, based on the processed data and precomputed dimensionality reduction embeddings provided in the original studies. UMAP representations and gene expression plots were generated using the datasets and annotations available within the portal. Cluster annotations were adapted for clarity and harmonized across datasets, when necessary, without modifying the original clustering or dimensionality reduction.

A publicly available cohort of patients was analyzed using data from the TCGA TARGET GTEx ^33^ repository (https://xenabrowser.net/datapages/?cohort=TCGA%20TARGET%20GTEx). The clinical dataset included information on sample type and primary site. Samples with undefined or indeterminate data were excluded from the corresponding analyses. Analysis included RNA-seq data from 1391 patients, comparing gene expression profiles between breast cancer tumor samples (Breast tumor tissue, n=1092) and samples of breast tissue adjacent to the tumor (Breast tumor-adjacent tissue, n=113). This dataset was used to assess RET expression levels across different breast cancer subtypes and to perform correlation analyses between RET expression and selected markers of interest.

#### Statistics & reproducibility

Data are presented as mean or median ± SEM or ± SD, as indicated. Normality was assessed using the Shapiro–Wilk test. Comparisons between two groups were performed using a two-tailed Student’s t test for normally distributed data or a Mann–Whitney test for non-normally distributed data. Comparisons among multiple groups were analyzed using one-way ANOVA followed by Tukey’s multiple-comparisons test for normally distributed data, or the Kruskal–Wallis test followed by Dunn’s multiple-comparisons test for non-normally distributed data. Outliers were identified using the ROUT method implemented in GraphPad Prism and excluded from the analysis when justified. Pearson or Spearman correlation analysis was performed to study correlation between variables. For all tests, differences were considered significant at a two-sided p < 0.05. p-values are indicated either on bar graphics or figure legends. For analysis of human material, covariates compared to tumor subtypes were tested by ANCOVA with Freedman-Lane permutation.

Sample size for *in vitro* experiments was determined based on experimental reproducibility and previous experience in similar assays. Sample size for animal experiments was determined based on previous experimental experience and in accordance with the principles of the 3Rs (Replacement, Reduction, and Refinement), aiming to minimize animal use while ensuring sufficient biological replicates for robust statistical analysis. The number of animals in each group is determined by the statistical power that is required to detect significant biologically relevant differences. A meaningful difference on means at least 80% power for one- and two-sided testing. Animals that presented disease or had been bitten because fighting in the cage were excluded. Animals were randomized into groups. Technicians and/or students were blinded by analyzing samples, except in some experiments where samples needed to be loaded correctly. All displayed points represented biological replicates. In figure legends, numbers of replicates (wells or individual animals) are indicated by n and the number of independent experiments is stated as N. All analyses were performed using Excel, R studio and GraphPad PRISM 8 software. The statistical details for all the experiments were indicated in the figure legends.

## ACKNOWLEDGEMENTS

This work was financially supported by Agencia Nacional de Promoción Científica y Tecnológica (AGENCIA, PCE-GSK-052 to A.G., J.P.F, M.dM., E.C.K, and O.A.C.; PICT-2019-02850 to A.G.; AGENCIA postdoctoral fellowship to A.L. and M.D.P), by Argentinian Research Council (CONICET Doctoral Fellowship to S.A.V., M.A.M, M.G.S., P.A.); by the the Federation of European Biochemical Societies, the Pan American Association for Biochemistry and Molecular Biology and the European Association for *Cancer Res*earch (IUBMB-FEBS-PABMB and EACR, respectively, to S.A.V for internships in G.S. laboratory), by Agencia Estatal de Investigación (AEI, PID2022-138525OB-I00 funded by MICIU/AEI/10.13039/501100011033 and by FEDER/EU funds to G.S.), by CRIS Contra el Cáncer (PR_EX_2024-22 to G.S.), and by the Instituto de Salud Carlos III (ISCIII, PMP21/00057, IMPACT-2021 Precision Medicine Infrastructure funded by ISCIII and the European Union through FEDER/FSE, Next Generation EU and the Plan de Recuperación, Transformación y Resiliencia to G.S.), by Margarita Salas and the Juan de la Cierva Program (CA1/RSUE/2021-00614 and FJC2020-043126-I, respectively, to G.S. to support A.C.A.), by CNIO Friends (postdoctoral fellowship to J.I.J-L). We thank the CNIO genomic and microscopy Unit for technical support and assistance with data analysis, as well as the staff of the Animal Facilities of both institutions for their help and support with the animal experiments. We are especially grateful to Dr. Nancy E. Hynes, who revealed the first evidence of RET-driven breast lesions that triggered our investigations. We thank Occident Foundation for strongly supporting the collaboration between Gattelli’s and Sabio’s laboratories.

## AUTHORS CONTRIBUTIONS

Conceptualization: S.A.V., G.S., A.G.; Methodology: S.A.V., A.L., I.R-G, I.N. M.D.P., P.A., A.A., L. L-V., M.L.-, E.R., A.G.; Bioinformatic analysis: S.A.V., M.G.S., M.dM., J.I.J.L.,; Validation: S.A.V., A.L.; Formal analysis: S.A.V., A.L., A.G.; Investigation: S.A.V., C.S.-L., E.C.K., G.S., A.G.; Resources: J.P.F., M.dM., E.C.K, G.S., A.G.; Clinical samples resources: D.M., D.F., N.J., E.Q., L.L., S.V., G.A., A.L., E.C.K., E.W. Data curation: M.G.S., M.A.M., M.dM. J.I.J.L.; Writing - original draft: S.A.V., G.S., A.G.; Writing - review & editing: S.A.V., C.S-L., O.A.C., G.S., A.G.; Visualization: S.A.V., A.L., M.A.M, G.N.H; Supervision: M.dM., G.S., A.G.; Project administration: A.G.; Funding acquisition: M.dM., J.P.F., O.A.C., E.C.K., G.S. A.G.

## SUPLEMMENTARY INFORMATION

Supplementary Information is available for this paper.

