## Supplementary figures & legends (Fig. S1-5) & Supplementary tables & footnotes (Table S1-4) for "TUMOR–PRE-ADIPOCYTE CROSSTALK SUSTAINS BREAST CANCER GROWTH VIA RET SIGNALLING"

SUPPLEMENTARY INFORMATION

Supplementary figures & legends (Fig. S1-5)

Supplementary tables & footnotes (Table S1-4)

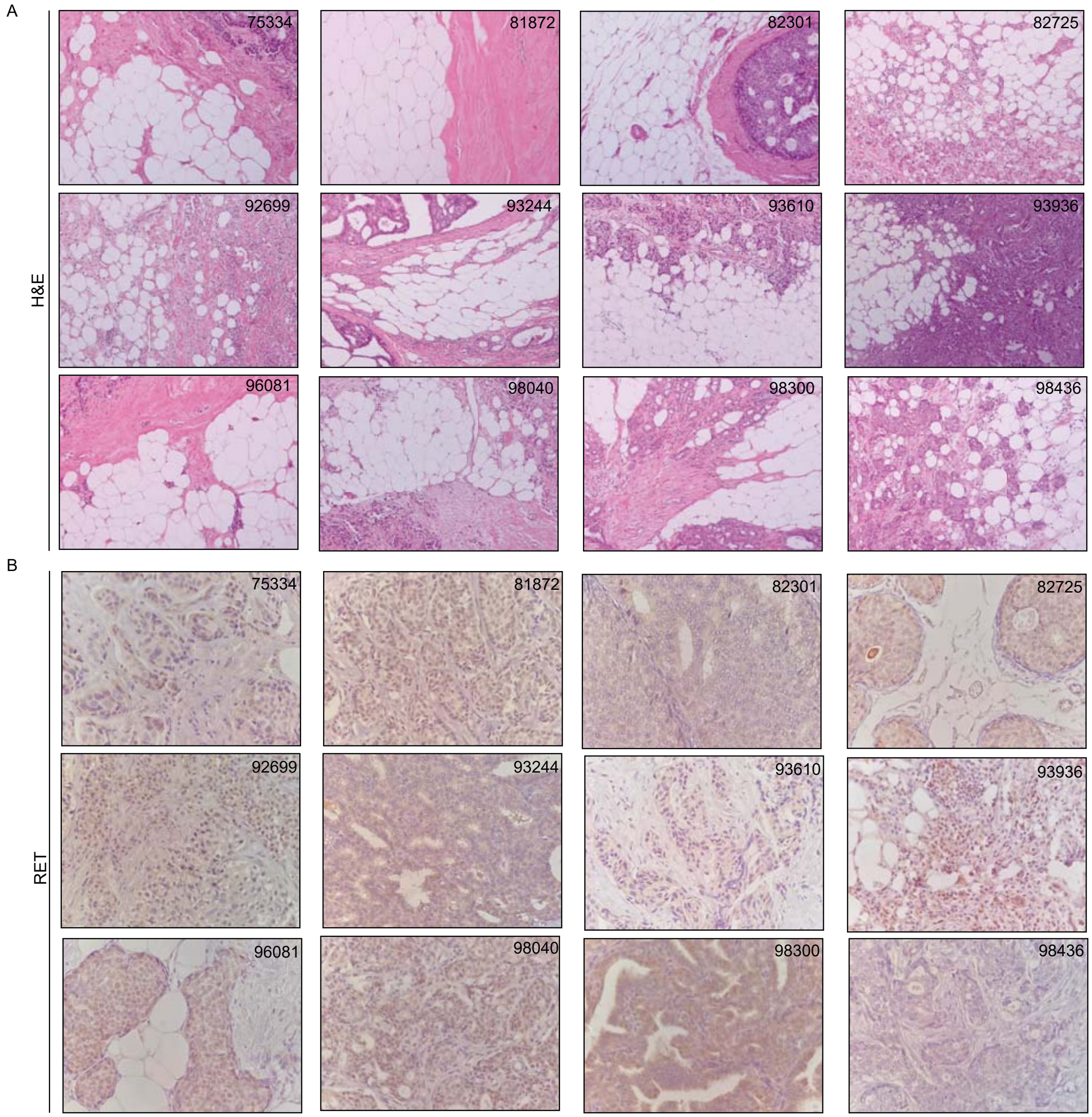
A
H&E
75334
81872
82301
82725
92699
93244
93610
93936
96081
98040
98300
98436
B
RET
75334
81872
82301
82725
92699
93244
93610
93936
96081
98040
98300
98436
Figure S1.

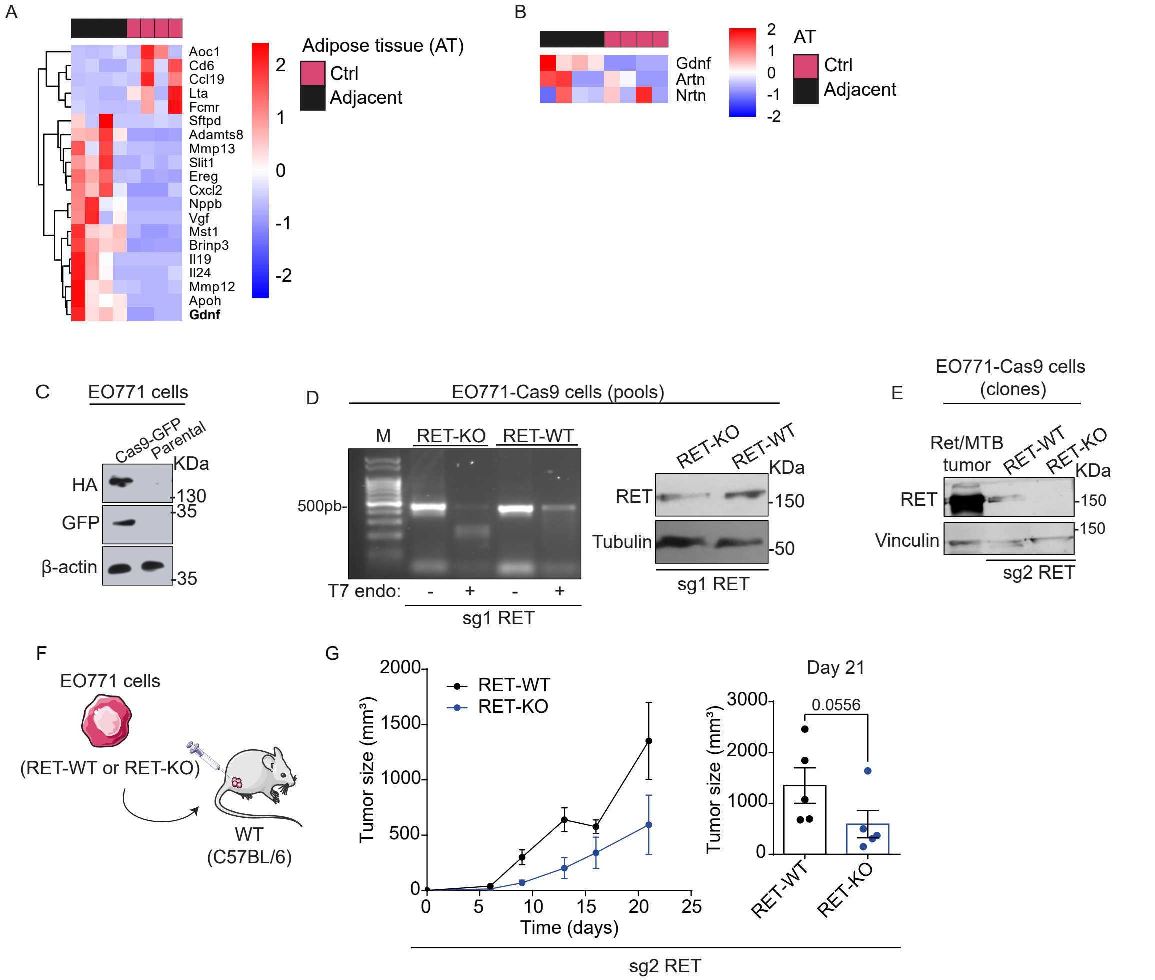

A
Adipose tissue (AT)
Ctrl
Adjacent
Aoc1
Cd6
Ccl19
Lta
Fcmr
Sftpd
Adamts8
Mmp13
Slit1
Ereg
Cxcl2
Nppb
Vgf
Mst1
Brinp3
Il19
Il24
Mmp12
Apoh
Gdnf
2
1
0
-1
-2
B
AT
Ctrl
Adjacent
Gdnf
Artn
Nrtn
C
EO771 cells
Cas9-GFP
Parental
KDa
HA
GFP
β-actin
130
35
35
D
EO771-Cas9 cells (pools)
M
RET-KO
RET-WT
500pb-
T7 endo: - + - +
sg1 RET
RET-KO
RET-WT
KDa
RET
Tubulin
150
50
sg1 RET
E
EO771-Cas9 cells (clones)
Ret/MTB tumor
RET-WT
RET-KO
KDa
RET
Vinculin
150
150
sg2 RET
F
EO771 cells
(RET-WT or RET-KO)
WT (C57BL/6)
G
RET-WT
RET-KO
Tumor size (mm³)
Time (days)
Day 21
0.0556
sg2 RET


Figure S2.

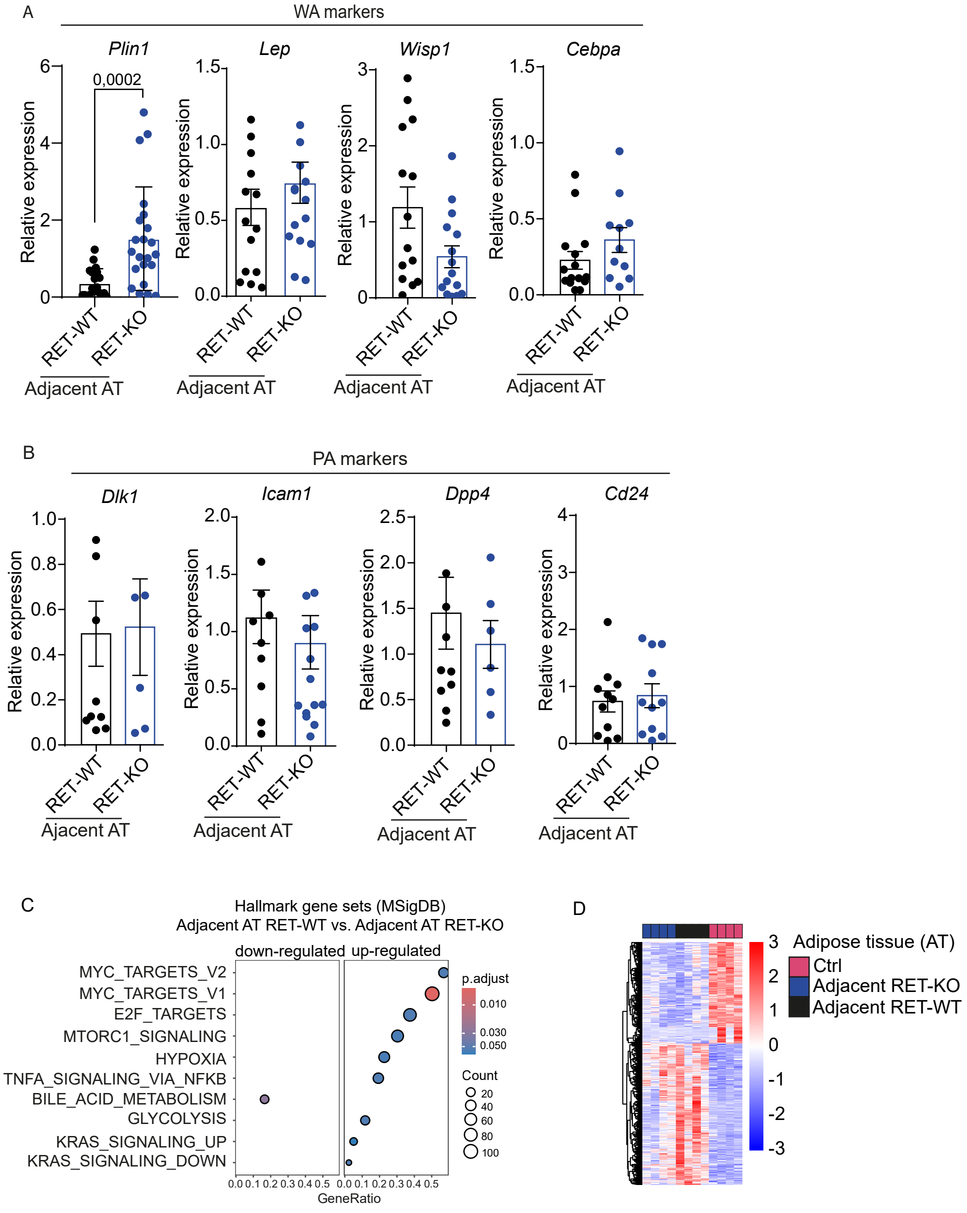

A
WA markers
Plin1
Lep
Wisp1
Cebpa
0,0002
Relative expression
RET-WT
RET-KO
Adjacent AT
B
PA markers
Dlk1
Icam1
Dpp4
Cd24
Ajacent AT
C
Hallmark gene sets (MSigDB)
Adjacent AT RET-WT vs. Adjacent AT RET-KO
down-regulated
up-regulated
MYC_TARGETS_V2
MYC_TARGETS_V1
E2F_TARGETS
MTORC1_SIGNALING
HYPOXIA
TNFA_SIGNALING_VIA_NFKB
BILE_ACID_METABOLISM
GLYCOLYSIS
KRAS_SIGNALING_UP
KRAS_SIGNALING_DOWN
p.adjust
0.010
0.030
0.050
Count
20
40
60
80
100
GeneRatio
D
Adipose tissue (AT)
Ctrl
Adjacent RET-KO
Adjacent RET-WT


Figure S3.

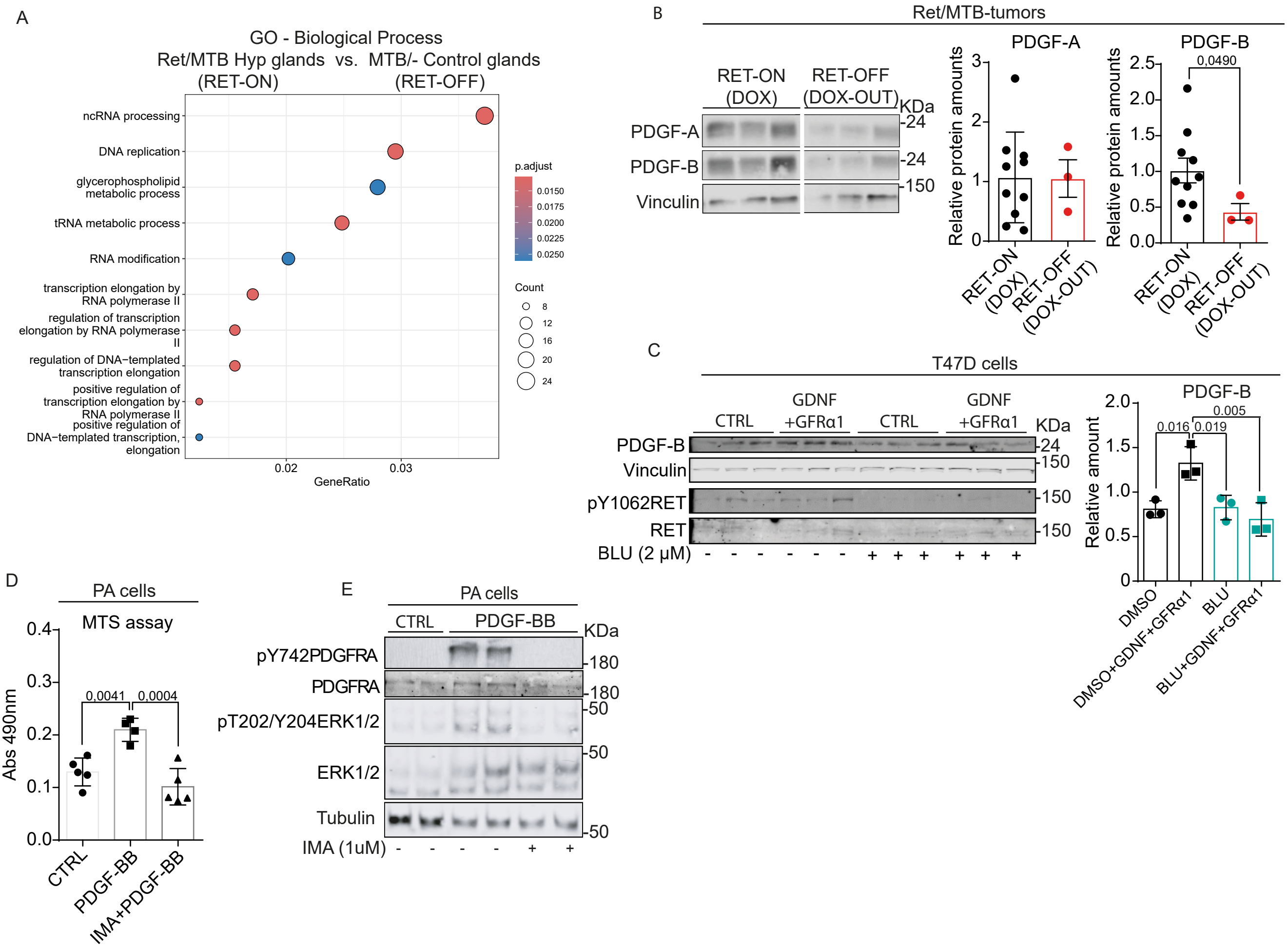
A
GO - Biological Process
Ret/MTB Hyp glands vs. MTB/- Control glands
(RET-ON) (RET-OFF)
ncRNA processing
DNA replication
glycerophospholipid metabolic process
tRNA metabolic process
RNA modification
transcription elongation by RNA polymerase II
regulation of transcription elongation by RNA polymerase II
regulation of DNA−templated transcription elongation
positive regulation of transcription elongation by RNA polymerase II
positive regulation of DNA−templated transcription, elongation
0.02
0.03
GeneRatio
p.adjust
0.0150
0.0175
0.0200
0.0225
0.0250
Count
8
12
16
20
24
B
Ret/MTB-tumors
RET-ON (DOX)
RET-OFF (DOX-OUT)
KDa
PDGF-A
PDGF-B
Vinculin
-24
-24
-150
PDGF-A
PDGF-B
Relative protein amounts
0,0490
RET-ON (DOX)
RET-OFF (DOX-OUT)
C
T47D cells
CTRL
GDNF +GFRα1
CTRL
GDNF +GFRα1
KDa
PDGF-B
Vinculin
pY1062RET
RET
BLU (2 μM)
-24
-150
-150
-150
PDGF-B
Relative amount
0.016
0.019
0.005
DMSO
DMSO+GDNF+GFRα1
BLU
BLU+GDNF+GFRα1
D
PA cells
MTS assay
Abs 490nm
0,0041
0,0004
CTRL
PDGF-BB
IMA+PDGF-BB
E
PA cells
CTRL
PDGF-BB
KDa
pY742PDGFRA
PDGFRA
pT202/Y204ERK1/2
ERK1/2
Tubulin
IMA (1uM)
-180
-180
-50
-50
-50

Figure S4.

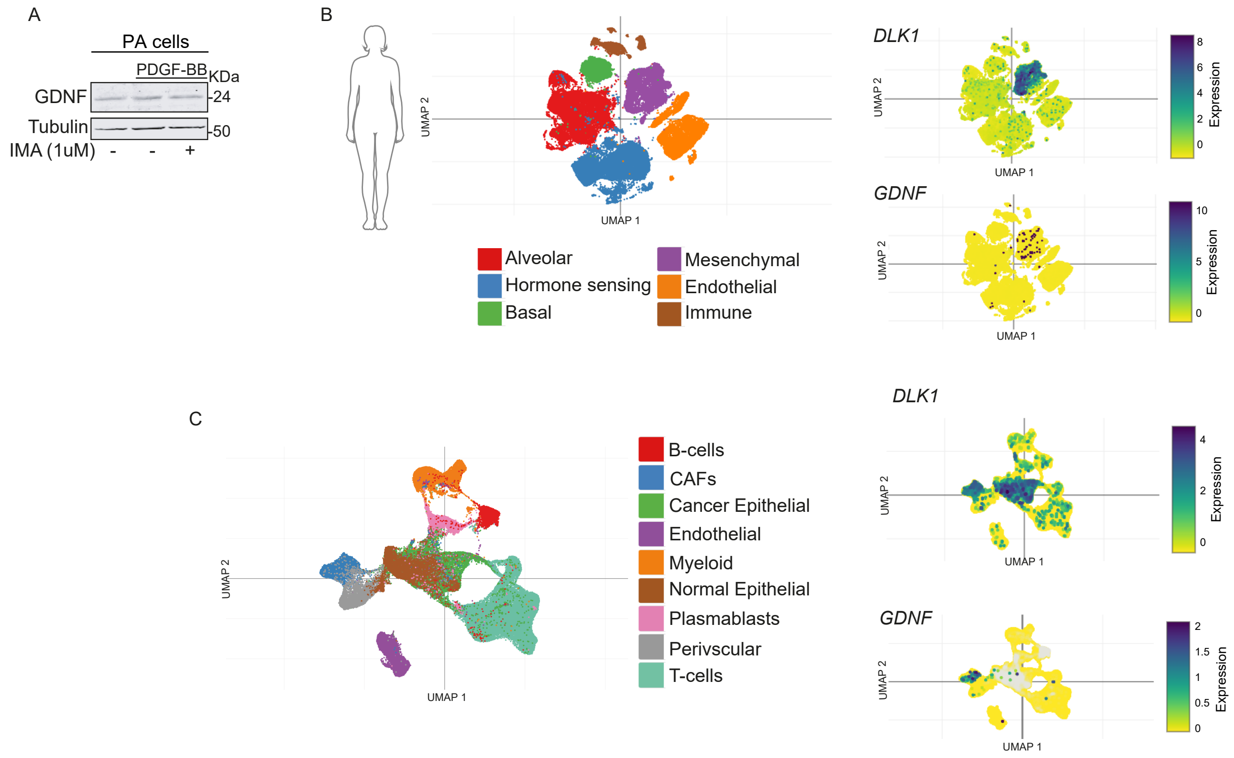
A
PA cells
PDGF-BB
KDa
GDNF
-24
Tubulin
-50
IMA (1uM) - - +
B
UMAP 2
UMAP 1
Alveolar
Hormone sensing
Basal
Mesenchymal
Endothelial
Immune
DLK1
Expression
8
6
4
2
0
GDNF
10
5
0
C
B-cells
CAFs
Cancer Epithelial
Endothelial
Myeloid
Normal Epithelial
Plasmablasts
Perivscular
T-cells
DLK1
4
2
0
GDNF
2
1.5
1
0.5
0

Figure S5.

***Figure S1. Staining in breast tumor tissue from patient cohort.***

(A) H&E- and (B) RET-staining in breast tumor tissue (Tumor) cores from patients (n=12). H&E stains display portion of adipose tissue next to the tumors. On pictures, the number of BC patient ID is indicated. Images were acquired at 10× or 40x magnification for H&E- or RET-staining respectively.

***Figure S2. RNA-seq analysis and generation of EO771 RET-KO cells and tumor allografts.***

(A-B) Related to Fig. 1. Heatmap of top 20 genes encoding soluble factors that arises from the analysis of the RNA-seq data performed in tumor-adjacent adipose tissue (Adjacent AT) respect to adipose tissue control (Ctrl AT) in the EO771 mouse mammary tumor model (n=4).

(C) Analysis of WB for GFP on lysates from EO771 Cas9-expressing cells (HA-GFP-Cas9) and the corresponding controls. Membrane was also blotted for HA tag protein as additional control.

(D) T7 endonuclease assay to confirm deletion efficiency was performed in DNA from EO771-Cas9 cells subject to edition. An agarose gel is showing the RET-specific PCR products corresponding to RET-WT and RET-KO cells treated or not with T7 enzyme. In parallel, the analysis of RET expression at protein level was assessed by WB in EO771 cell pools lysates (sg1RET).

(E) Analysis of RET expression at protein level by WB in RET-WT and RET-KO EO771 cell clones (sg2RET). Lysates of Ret/MTB-tumor were used as positive controls of RET expression.

(F) Scheme of RET-KO EO771-tumor allograft model. RET-WT or RET-KO EO771 tumor cell lines (sg2 RET) were injected into female mice to generate EO771-derived tumors.

(G) *In vivo* tumor growth of RET-WT or RET-KO EO771-tumor allografts (sg2RET). Tumor volumes were measured after mammary fat pad injection of EO771 into female mice (RET-WT, n=5; RET-KO, n= 5). Data are presented as mean ± SEM. Each dot in the curve represents the mean of a group of animals. In the bar chart, each dot represents an animal. Mann-Whitney test was used; p=0.0556.

***Figure S3. Analysis of adipocyte differentiation markers in EO771-tumor model.***

(A-B) Related to Fig. 3. Analysis of expression of specific adipocyte differentiation markers were performed by RT-qPCR on samples of Adjacent AT from EO771 RET-WT or RET-KO EO771-tumor bearing animals. Expression levels for (A) WA markers (*Plin1*, *Lep*, *Wisp1, Cebpa*) and (B) PA or adipocyte precursor cell markers (*Dlk1*, *Pdgfra*, *Icam1, Dpp4*, *Cd24*) are indicated. Each dot represents an individual animal (n=6-21) from 3 independent experiments. Data are presented as mean ± SEM. Unpaired Student's t or Mann-Whitney test was used; p values are indicated on bar graphics.

(C-D) Hallmark gene set analysis (MSigDB) of RNA-seq data from tumor-adjacent adipose tissue (Adjacent AT) from RET-WT versus RET-KO tumors in the EO771 mouse mammary tumor model from independent animals (n=4). GO enrichment significance was assessed using Fisher's exact test, with p-values corrected for multiple testing using the Benjamini–Hochberg false discovery rate (FDR). Heatmap shows the distribution of gene expression between groups.

***Figure S4. PDGF-B is a RET-induced factor which increases viability of pre-adipocytes (PA) cells.***

(A) Analysis of GO Biological Process of the RNA-seq data performed in the RET-overexpressing Ret/MTB bitransgenic glands (RET-ON) respect to MTB/- control glands (RET-OFF) from independent animals (n=4). GO enrichment significance was assessed using Fisher's exact test, with p-values corrected for multiple testing using the Benjamini–Hochberg false discovery rate (FDR).

(B) PDGFs levels were confirmed in tumor tissue from the RET/MTB-tumor mouse model. Results from 3 independent animals WB analysis are representatives. Corresponding quantification is shown. Each dot represents an individual animal (n=3-10, 2 independent experiments). Unpaired Student's t test was used; *p=0.0490.

(C) Related to Fig. 4. T47D cultures were stimulated with GDNF and co-receptor (GFR$\alpha$1; 100 ng/ml) and subsequently, PDGFs ligands were analyzed in lysates by WB. Results are representatives of at least 3 independent experiments. Corresponding quantification is shown. Each dot represents an individual well (n=3). One-way ANOVA test was used followed by Tukey's multiple comparation test p-values are indicated on bar graphics.

(D) MTS assay was performed to assess viability in PA-treated cultures. PDGF-BB (0,1 mg/ml) was directly added to PA cell cultures previously treated or not with IMA (1$\mu$M) and cell viability was assessed 48 hours later. Each dot represents an individual well (n=5) for 4 independent experiments (N=4). One-way ANOVA test was used followed by Tukey's multiple comparation test; p-values are indicated on bar graphics.

(E) Control of levels of activation/inhibition of PDGFR and ERK1/2 by PDGF-BB (0,1mg/ml) ± IMA (1mM) is shown by WB analysis.

***Figure S5. Pre-adipocyte (PA)-like cells analysis for GDNF expression.***

(A) Expression levels of RET ligand (GDNF) were analyzed in PA cultures treated or not with PDGF-BB (0,1 mg/ml) in absence or presence of PDGFR inhibitor (IMA, 1$\mu$M).

(B) Analysis of published single-cell RNA-seq data visualized using Uniform Manifold Approximation and Projection (UMAP) for human normal breast tissue (n=16)[50]. The left panel shows the separation of major cell-type populations, indicated by distinct colors. The right panels display *DLK1* and *GDNF* expression levels, respectively, across the identified cell populations.

(C) Analysis of published single-cell RNA-seq data visualized using UMAP for human breast tumors (n=26) [49]. The left panel shows the separation of major cell-type populations, indicated by distinct colors. The right panels display *DLK1* and *GDNF* expression levels, respectively, across the identified cell populations.

***Table S1. Histopathological features of the patient cohort.***

| **BC patient** (ID) | **Molecular classification** (by evaluation of IHC) | **Age** (years) | **Histological grade** (by Nottingham score) | **Ki-67** (% by evaluation of IHC) | **Used for** (technique) |
|---|---|---|---|---|---|
| 75334 | ER+ PR+ HER2- | 59 | G2 | 20 | H&E, IHC, RT-qPCR |
| 81872 | ER+ PR+ HER2- | 57 | G3 | 15 | H&E, IHC, RT-qPCR |
| 82301 | ER+ PR+ HER2- | 45 | G3 | 35 | H&E, IHC |
| 82725 | ER+ PR+ HER2- | 62 | G2 | 10 | H&E, IHC, RT-qPCR |
| 92699 | ER- PR- HER2- | 58 | G3 | 70 | H&E, IHC |
| 93244 | ER+ PR+ HER2- | 51 | N/A | 5 | H&E, IHC |
| 93610 | ER+ PR+ HER2+ | 68 | G2 | 60 | H&E, IHC |
| 93936 | ER- PR- HER2+ | 58 | G3 | 60 | H&E, IHC |
| 96081 | ER+ PR+ HER2- | 75 | G2 | 7 | H&E, IHC, RT-qPCR |
| 98040 | ER+ PR+ HER2+ | 78 | G2 | 30 | H&E, IHC |
| 98300 | ER+ PR+ | 61 | N/A | N/A | H&E, IHC, RT-qPCR |
| 98436 | ER+ PR+ HER2- | 79 | G2 | 30 | H&E, IHC, RT-qPCR |
| 94914 | ER+ PR- HER- | 64 | G3 | 5 | RT-qPCR |
| 74497 | ER+ PR- HER- | 72 | G2 | 15 | RT-qPCR |
| 78002 | N/A | N/A | N/A | N/A | RT-qPCR |
| T56 | N/A | N/A | N/A | N/A | RT-qPCR |
| 96641 | ER+ PR+ HER- | 49 | N/A | 10 | RT-qPCR |
| 97251 | N/A | 54 | G2 | N/A | RT-qPCR |
| 81919 | ER+ PR+ HER- | 43 | G3 | 60 | RT-qPCR |

Clinical and histopathological parameters of biopsies from female patients with breast cancer who underwent surgery (2016-2020, n=19) at the Marie Curie Municipal Oncology Hospital in Buenos Aires, Argentina. Molecular classification of tumor subtype is indicated by IHC and posterior pathological evaluation. ER: estrogen receptor, PR: progesterone receptor, HER2: Her2/ErbB2 receptor amplification, Ki-67: proliferation marker; Patient age, is indicated in years; G1-G3: tumor grade by Nottingham Histologic Score; N/A: not available. For each BC patient's ID, the assays performed in this study are indicated.

***Table S2. Analysis of histological images of each sample from the patient cohort.***

| **BC patient** | **Adipose stromal tissue** | | | | **Tumor lesion** |
|---|---|---|---|---|---|
| | **Mean adipocyte area** ($\mu m^2$) | | **Adipocyte count** (cell number) | | **Mean RET intensity** (a.u.) |
| (ID) | Distant AT | Adjacent AT | Distant AT | Adjacent AT | |
| 75334 | 2515.01 | 2368.69 | 1277 | 1776 | 83.26 |
| 81872 | 3053.45 | 2242.36 | 1698 | 2276 | 86.85 |
| 82301 | 2316.99 | 1604.76 | 1615 | 1690 | 90.51 |
| 82725 | 1991.50 | 1317.63 | 483 | 1636 | 78.24 |
| 92699 | 1735.08 | 1458.21 | 310 | 1158 | 92.17 |
| 93244 | 2851.80 | 2273.78 | 922 | 1306 | 103,34 |
| 93610 | 882.15 | 764.86 | 249 | 1322 | 68.84 |
| 93936 | 776,00 | 759.79 | 614 | 1033 | 62.91 |
| 96081 | 2968.19 | 1984.30 | 544 | 1963 | 95.13 |
| 98040 | 1447.99 | 966.60 | 955 | 1416 | 89.05 |
| 98300 | 2726.32 | 1406.18 | 651 | 1798 | 10965 |
| 98436 | 2641.53 | 2011.47 | 738 | 2076 | 64.625 |

'Distant' adipocytes (>2000 µm away from tumor lesion) and 'Adjacent' adipocytes (within 2000 µm surrounding tumor lesion) are analyzed. The parameters measured, the values obtained for adipocyte counting (cell number) and the mean area of adipocytes ($\mu m^2$) in the AT are displayed. The mean intensity of RET-positive cells stained in arbitrary units (a.u.) in the tumor is indicated.

***Table S3. RET expression positively correlates with markers of immature adipocytes in the tumor adjacent tissue of breast cancer patients.***

| Correlation coefficient (-1 0 1) | | RET | |
|---|---|---|---|
| | | Breast tumor-adjacent tissue (n = 113) | Breast tumor tissue (n = 1092) |
| White adipocytes (WA) | *ADIPOQ* | -0,11 ns[b] | 0,06 *[b] |
| | *PPARG* | -0,15 ns[b] | -0,07 *[b] |
| | *LEP* | -0,24 **[b] | -0,02 ns[b] |
| Pre- adipocytes (PA) | *DLK1* | 0,39 ***[b] | -0,04 ns[b] |
| | *PDGFRA* | -0,07 ns[b] | -0,14 ****[b] |
| | *ICAM1* | -0,11 ns[a] | -0,31 ****[b] |

Correlation analysis of adipocyte differentiation marker genes (*ADIPOQ*, *PPARG*, *LEP* for WA; *DLK1*, *PDGFRA*, *ICAM1* for PA) and RET expression in human breast biopsies (TCGA TARGET GTEx) corresponding to breast tumor-adjacent tissue (n=113) or tumor tissue (n=1092). WA: White adipocytes markers; PA: Pre-adipocytes markers. Pearson (a) or Spearman (b) correlation test was used; ns: not significant; *p<0.05; **p<0.005; ***p<0.001; ****p<0.0001.

***Table S4. RET-expressing breast tumors correlate with PDGF-B ligand in tumor tissue in patient samples.***

| Correlation coefficient (-1 0 1) | RET | |
| --- | --- | --- |
| | Breast tumor-adjacent tissue (n = 113) | Breast tumor tissue (n = 1092) |
| *PDGFA* | 0,13 | -0,06 |
| | ns[b] | ns[b] |
| *PDGFB* | -0,09 | 0,2 |
| | ns[b] | ****[b] |

Correlation analysis of expression of PDGF ligands (*PDGFA*, *PDGFB*) and RET expression in human breast tumor (n=1092) and adjacent tissue (n=113) biopsies (TCGA TARGET GTEx). Spearman (a) or Pearson (b) correlation test was used; ns: not significant; *$p<0.05$; **$p<0.01$; ***$p<0.001$; ****$p<0.0001$.
